# Somatic-to-germline transmission of horizontally acquired extrachromosomal circular DNA in Brassica graft chimeras

**DOI:** 10.64898/2026.05.08.723653

**Authors:** Aijun Zhang, Han Zhou, Wanmu Zhou, Kexin Xu, Tingjin Wang, Xiaoyang Wan, Junxing Li, Lu Yuan, Li Huang, Yanhong Zhou, Susheng Gan, Yongfeng Guo, Qinjie Chu, Yan Wang, Ningning Yu, Hannah Rae Thomas, Ke Liu, Liping Chen

## Abstract

Whether somatically acquired traits can contribute to heritable variation has remained an open question in plant biology for over a century. Here, using graft-induced periclinal chimeras between *Brassica juncea* and *Brassica oleracea*, we identified extrachromosomal circular DNA (eccDNA) as mobile genetic elements capable of crossing histological boundaries and entering the germline. We demonstrated that grafting could promote the horizontal transfer of eccDNA between somatic cell layers, enabling its stable maintenance in recipient tissues and permitting transmission through sexual reproduction across multiple generations. We found that transmitted eccDNA is non-randomly distributed across the genome, preferentially originating from gene-dense regions, and enriched for features associated with molecular persistence, such as inverted repeats, hairpin-forming sequences, and autonomously replicating sequence (ARS) consensus motifs. These genetic regions conferred replication competence, as evidenced by autonomous propagation in an ARS-less heterologous yeast system. Sexual progeny carrying graft-acquired eccDNA exhibited reproducible and lineage-dependent alterations in leaf morphology and drought tolerance, accompanied by coordinated transcriptional reprogramming. Notably, a subset of inherited eccDNA remained transcriptionally active in progeny, producing transcripts absent from self-grafted controls. Our findings establish eccDNA as heritable extrachromosomal elements that link graft-mediated somatic genetic transfer with stable germline transmission, thereby expanding the molecular scope of heritable variation in plants and providing a conceptual framework for graft-based trait transmission.

## Introduction

Whether traits acquired in somatic tissues can be transmitted across generations has remained an unresolved question in genetics. Classical genetic theory holds that heritable information flows exclusively through the germline, whereas somatic variations, although continuously shaped by environmental cues and biotic interactions, are generally considered evolutionarily transient and excluded from inheritance ^1^. Although early theoretical models speculated that somatically derived factors might influence offspring phenotypes ^2,3^, direct experimental evidence supporting such transmission has remained limited. In plants, it therefore remains unclear whether specific molecules acquired during somatic variation can be transmitted across tissues, reach the germline, and ultimately contribute to heritable variation.

Grafting between different species is an important form of biotic interaction within the plant kingdom ^4–6^. As a classic form of somatic communication, grafted rootstocks can induce phenotypic changes in the scion ^7–10^. However, whether graft-induced molecular changes can be transmitted to sexual progeny remains poorly understood. Progress has been hindered by two key limitations: the lack of experimental systems that clearly distinguish and trace somatic and germline lineages, and the absence of a molecular vehicle capable of mediating cross-lineage transmission. Shoot apical meristem (SAM) grafting provides a framework to overcome these limitations. SAM grafting utilizes wounding to induce chimeric shoot formation at the graft site. The SAM is organized into three histological layers (L1, L2, and L3), with cells from the L2 layer giving rise to gametes and contributing directly to sexual progeny ^11,12^. Periclinal chimeras generated by SAM grafting, which consist of stable, genetically distinct cell layers within a single individual, provide a unique opportunity to test whether molecules originating in somatic layers can traverse tissue boundaries and access the reproductive lineage ^13–16^.

Among potential carriers of graft-derived inheritance, extrachromosomal circular DNA (eccDNA) is an intriguing class of molecules ^17–19^. EccDNA molecules are circular DNA excised from chromosomal regions and widespread across eukaryotes ^20–22^. EccDNA is structurally diverse, physically mobile, and accumulates dynamically in response to stress or development ^20–25^. Recent work has demonstrated that eccDNA can move horizontally from grafted rootstocks and remain stably present in somatic scion tissues and their clones, influencing growth-related traits ^26^. However, whether somatically acquired eccDNA during grafting can cross the soma– germline boundary and affect sexual progeny remained unknown.

The molecular basis by which acquired eccDNA might persist through cell divisions and escape dilution during development remains an open question. In eukaryotic systems, the maintenance of eccDNA is often associated with the presence of cis-acting replication elements. In yeast, autonomously replicating sequences (ARSs) consist of short DNA segments sufficient to support plasmid replication independent of the chromosomes ^27^. Canonical ARSs typically span 100–200 bp and contain an AT-rich 11-bp autonomously replicating sequence consensus (ACS) that recruits the origin recognition complex (ORC), while auxiliary elements enhance origin efficiency by stabilizing ORC association, facilitating replicative helicase (MCM) loading, and shaping local chromatin architecture ^28–31^.

Although replication origins in higher plants remain less well defined, accumulating evidence suggests that plant eccDNA can acquire autonomous or semi-autonomous replication capacity. In *Amaranthus palmeri*, eccDNA carrying herbicide-resistance genes form complex circular replicons that undergo rapid amplification under selective pressure, and some harbor ACS-like elements capable of driving replication in heterologous systems ^32–34^. In addition, inverted repeats, hairpin-forming sequences, and other secondary structures are known to facilitate replication and the stability of circular DNA in viruses and microbes ^29,35–37^. However, the contribution of secondary structures to eccDNA persistence in plants remains largely unexplored. Together, these observations suggest that at least a subset of plant eccDNAs may encode intrinsic signals that enable their persistence, amplification and inheritance, rather than being passively carried during cell division.

Here, we addressed whether eccDNA, acquired through somatic exchange, can access the reproductive lineage and be propagated across generations. Using artificial Brassica periclinal chimeras generated between *B. juncea* and *B. oleracea* ^38,39^, we exploited the layered organization of the SAM to disentangle somatic and germline contributions. By regenerating individual cell layers and subsequently self-pollinating, or by directly selfing chimeric plants, we obtained sexual progeny in which graft-derived molecular signatures could be systematically assessed. Circle-seq profiling revealed extensive eccDNA transfer from somatic layers into the germline, with transmitted eccDNA enriched for hairpins, inverted repeats, and 11-bp ACS-like motifs. Functional assays in an ARS-deficient yeast system demonstrated that eccDNA-derived sequences are sufficient to support autonomous plasmid replication, providing direct evidence for their intrinsic role in replication. Independent molecular validation confirmed eccDNA transmission, and progeny carrying transferred eccDNA exhibited reproducible, lineage-dependent changes in leaf morphology and drought tolerance, accompanied by coordinated transcriptional reprogramming. Consistent with their heritable maintenance, a subset of transmitted eccDNA remained transcriptionally active in the progeny and produced transcripts that were undetectable in self-grafted controls. Together, our findings identify eccDNA-mediated transmission as a heritable extrachromosomal genetic process that links graft-induced somatic interactions with stable inheritance through the germline, thereby expanding the molecular scope of heritable variation in plants. This work establishes a conceptual and experimental basis for using graft-transmissible genetic elements as a source of heritable variation relevant to plant adaptation and crop improvement.

## Results

### Periclinal chimeras establish a traceable somatic–germline interface

To determine whether eccDNA acquired in somatic tissue through grafting can traverse histological layers into sexual reproduction, we established two periclinal chimeras between tuber mustard (*B. juncea;* T) and red cabbage (*B. oleracea*; C) using shoot apical meristem (SAM) grafting, following established protocols ^38^. The parents were designated TTT (*B. juncea*) and CCC (*B. oleracea*), and corresponding self-grafted (g) controls were generated for each genotype (g-TTT and g-CCC). Regeneration (r) from the L1 layer of the self-grafted controls produced clonal asexual progeny (rg-TTT and rg-CCC) ^16^, whereas self-pollination of these plants yielded sexual progeny (p) derived from the L2 layer (rgp-TTT and rgp-CCC), providing baseline references for somatic and sexual inheritance.

SAM grafting between TTT and CCC reproducibly generated two stable periclinal chimera types, termed TCC and TTC (L1-L2-L3) ^40^, with distinct L1-L3 layer compositions. In the TCC chimera, the outermost L1 layer was derived from *B. juncea*, whereas the inner L2–L3 layers originated from *B. oleracea*. In contrast, the TTC chimera consisted of L1–L2 layers from *B. juncea* and an L3 layer from *B. oleracea* (Fig. 1). This layered organization enabled heterologous eccDNAs acquired in somatic tissues to be spatially resolved and experimentally traced.

**Fig. 1.**
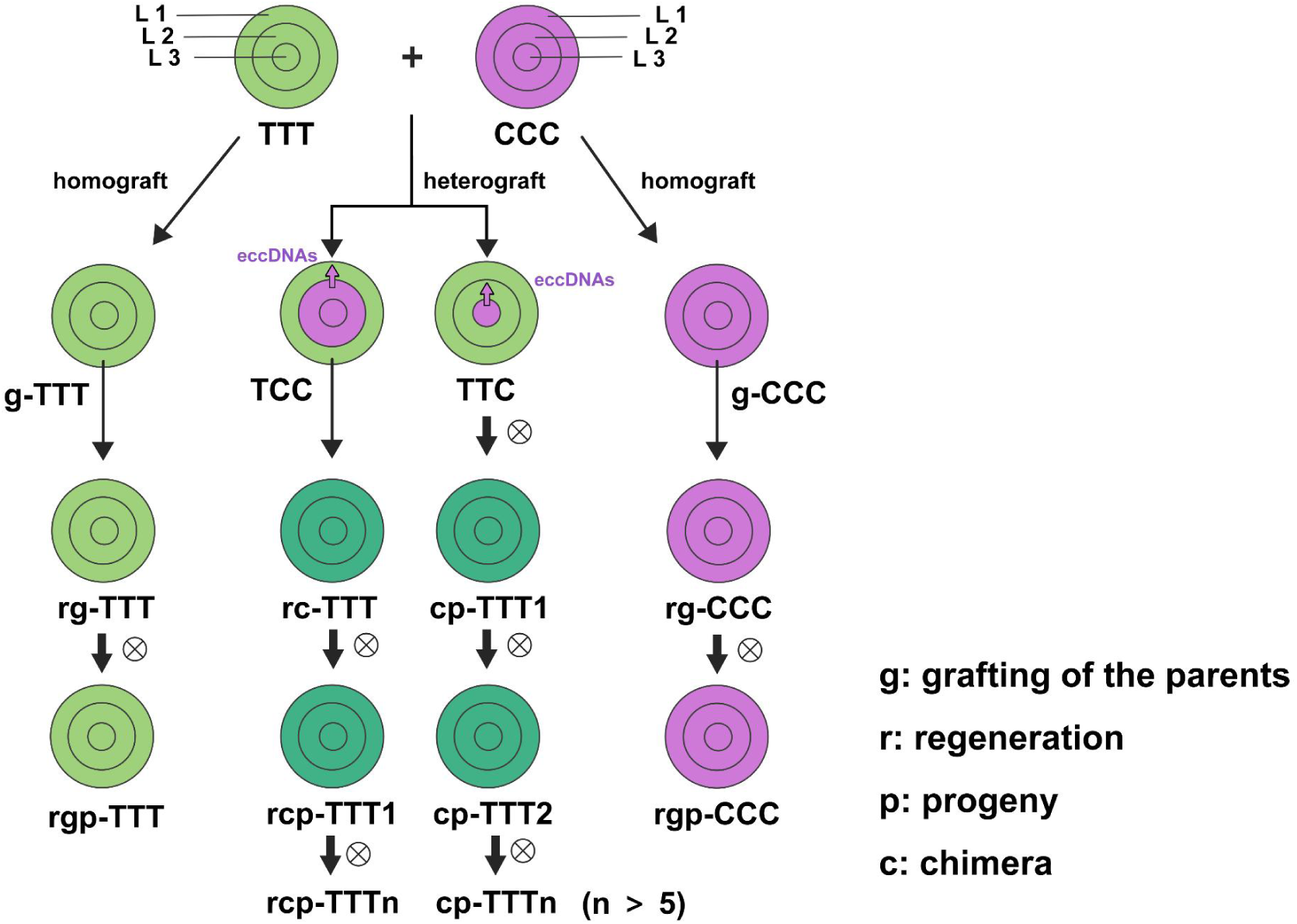
Generation of periclinal chimeras between *B. oleracea* and *B. juncea* and their derived *B. juncea*-lineage sexual progeny. *In vitro* heterografting between tuber mustard (*B. juncea*, TTT) and red cabbage (*B. oleracea*, CCC) produced periclinal chimeras TCC (L1=T; L2–L3=C) and TTC (L1–L2=T; L3=C). The L1 (T) layer of TCC was regenerated to generate rc-TTT plants, which were selfed to obtain rcp-TTT1 and subsequent progeny (rcp-TTTn). TTC chimeras were selfed directly to obtain cp-TTT1, and selfing of cp-TTT1 generated cp-TTT2 and additional progeny (cp-TTTn; n>5). Self-grafted parents (g-TTT and g-CCC) and their L1-regenerated controls (rg-TTT and rg-CCC) were selfed to produce rgp-TTT and rgp-CCC.

Exploiting this system, we obtained both asexual and sexual progeny with known lineage origins. To test whether graft-acquired eccDNAs present in the chimera can be carried through regeneration and subsequently detected in sexual offspring, regeneration from the L1 layer of the TCC chimera (c) produced clonal asexual progeny (rc-TTT), which were subsequently self-pollinated to generate sexual offspring (rcp-TTT). In parallel, to assess whether graft-acquired molecules can access the L2-derived reproductive lineage, TTC chimeras were self-pollinated, yielding sexual progeny designated cp-TTT. Because germ cells in Brassica are derived from the L2 layer, these sexual progenies provide a direct readout of whether genetic material acquired in somatic layers through grafting can cross tissue boundaries and enter the germline. Two representative progenies, rcp-TTT1 and cp-TTT2, were selected for detailed eccDNA profiling, phenotypic characterization and transcriptomic analyses (Fig. 1). Together, this experimental framework establishes lineage-resolved routes from chimeric tissues to L2-derived sexual progeny and enables the fate of graft-acquired eccDNAs to be followed and compared across progeny types.

### The eccDNA landscape across sexual progeny of Brassica chimeras

To characterize eccDNA acquired through SAM grafting and retained in sexual reproduction, we surveyed eccDNA six leaf samples. Circle-seq ^22,41^ was conducted on leaves from the parent species (*B. juncea* and *B. oleracea*), their self-grafted sexual progeny (rgp-TTT and rgp-CCC), and the sexual progeny derived from periclinal chimeras generated via regeneration (rcp-TTT1) or direct selfing (cp-TTT2) (Fig. 1, Fig. 2a). Two biological replicates were analyzed for each tissue type, yielding a total of twelve libraries. Following removal of chromosomal DNA contamination, eccDNA was enriched and amplified by random rolling circle amplification (rRCA), generating approximately 2.13 Gb of high-quality paired-end reads (Extended Data Table 1) suitable for genome-wide eccDNA profiling.

**Fig. 2.**
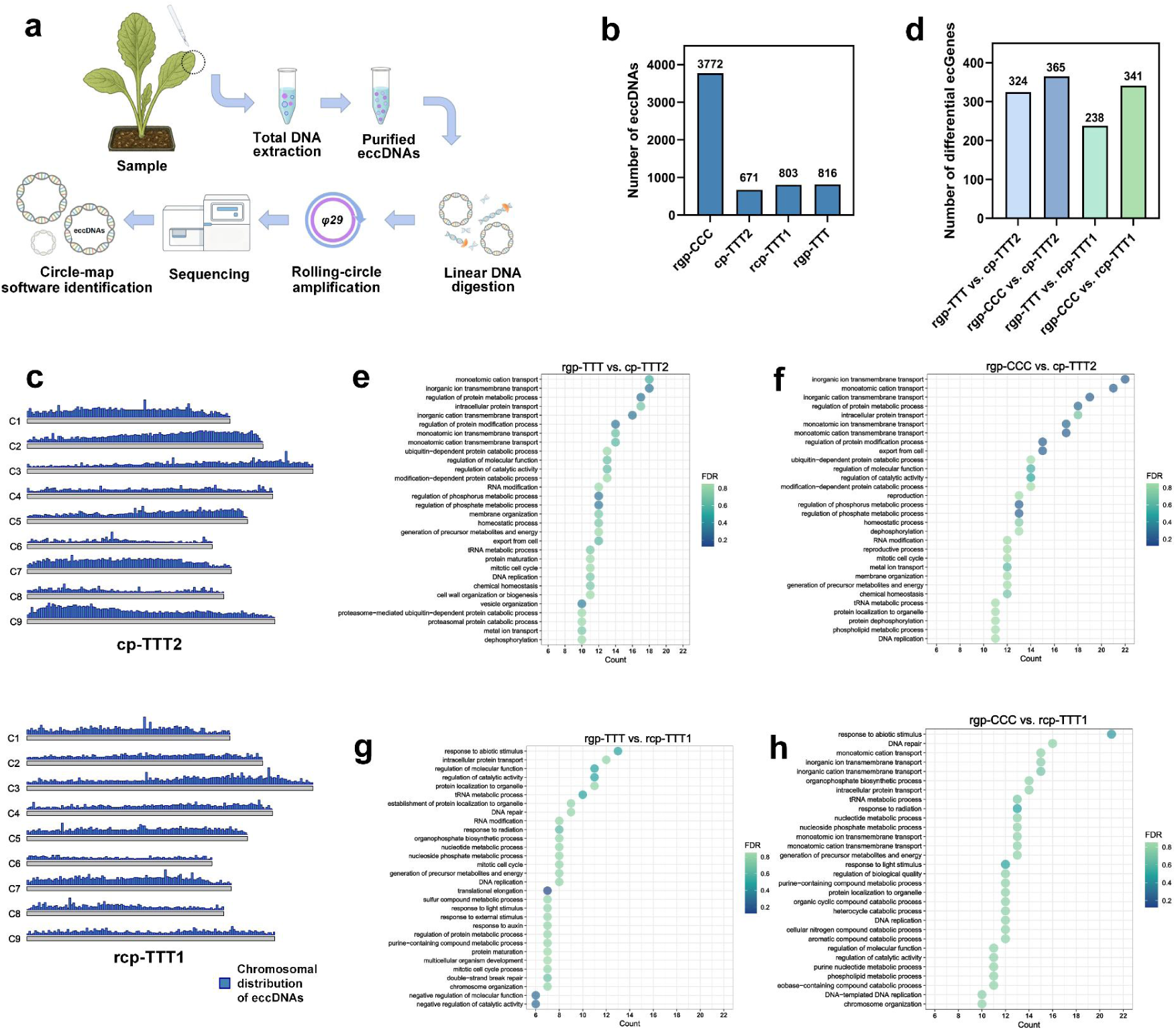
Global landscape of *B. oleracea*–derived eccDNAs and functional annotation of eccDNA-overlapping genes. **a,** Schematic overview of the Circle-seq workflow and Circle-Map calling used for eccDNA detection. **b,** Number of eccDNAs mapped to the *B. oleracea* reference genome in rgp-CCC, cp-TTT2, rcp-TTT1 and rgp-TTT. **c,** Karyoplots displaying the chromosomal distribution (C1-C9) of *B. oleracea* mapped eccDNAs identified in cp-TTT2 and rcp-TTT1. **d,** Number of differential ecGenes identified in pairwise comparisons indicated. ecGenes were defined as genes overlapped by eccDNA loci.. **e–h,** Gene Ontology (GO) biological process enrichment for differential ecGenes in rgp-TTT vs. cp-TTT2 (e), rgp-CCC vs. cp-TTT2 (f), rgp-TTT vs. rcp-TTT1 (g), and rgp-CCC vs. rcp-TTT1 (h). Dot size indicates gene count and colour indicates FDR.

All reads were uniquely aligned to the *B. oleracea* (CCC) reference genome, thereby quantifying *B. oleracea*–mapped (heterologous) eccDNAs across the six sample types (Extended Data Tables 2 and 3). Both the number and abundance of eccDNA varied markedly across these materials. The self-grafted cabbage progeny (rgp-CCC) exhibited the highest number of eccDNA (3,772), whereas the self-grafted mustard progeny (rgp-TTT) harbored substantially fewer (816) that could be aligned to the *B. oleracea* reference genome (Fig. 2b). Sexual progeny derived from periclinal chimeras displayed intermediate numbers of *B. oleracea*–mapped eccDNAs, consistent with retention of heterologous eccDNAs in the chimera-derived progeny (Fig. 2c and Extended Data Table 3). Chromosomal distribution analysis further revealed that eccDNA was broadly dispersed across the genome rather than confined to specific loci or chromosomes (Fig. 2c; Extended Data Fig. 1).

To further investigate the relationship between eccDNA aligned to the *B. oleracea* reference genome and annotated genes, we focused on eccDNA overlapping gene bodies (hereafter referred to as ecGenes) and compared their enrichment across different genetic backgrounds. Four pairwise comparisons were performed: cp-TTT2 versus rgp-TTT, cp-TTT2 versus rgp-CCC, rcp-TTT1 versus rgp-TTT and rcp-TTT1 versus rgp-CCC (Fig. 2d). Differentially enriched ecGenes was identified based on normalized junction read counts using edgeR (fold change ≥ 2.0, FDR ≤ 0.05), followed by gene-level annotation, and enrichment analysis ^42^.

Relative to rgp-TTT and rgp-CCC, rcp-TTT1 showed 238 and 341 differentially enriched ecGenes, respectively (Fig. 2d). Gene Ontology (GO) analysis revealed consistent enrichment of DNA replication and DNA repair–related processes, irrespective of the control background used (Fig. 2g, h). This convergence suggests that eccDNA present in rcp-TTT1 may be associated with genome maintenance and structural remodeling. When compared with rgp-TTT, additional enrichment of cell cycle regulation and auxin-responsive pathways were observed (Fig. 2g), consistent with previous reports implicating eccDNAs in developmental and hormonal regulation ^20^. By contrast, comparison with rgp-CCC highlighted enrichment of metabolic and catabolic processes (Fig. 2h), indicating genotype-dependent metabolic adjustment. Together, these patterns suggest that rcp-TTT1 carries eccDNA-associated functional signatures consistent with its phenotypic traits.

A comparable landscape was observed in cp-TTT2, which exhibited 324 and 365 differentially enriched ecGenes relative to rgp-TTT and rgp-CCC, respectively (Fig. 2d). In both comparisons, ecGenes were consistently enriched for DNA replication and mitotic cell cycle processes, alongside pathways related to ion transmembrane transport, protein modification and cellular homeostasis (Fig. 2e, f). The overlap in enriched categories with rcp-TTT1 suggests a shared eccDNA-associated response in chimera-derived progeny. Collectively, these results establish a shared eccDNA landscape in chimera-derived sexual progeny, providing a genomic framework for subsequent analyses of eccDNA transmission, inheritance, and functional consequences.

### Heterologous somatically acquired eccDNAs traverse the somatic–germline boundary

To directly test whether eccDNA acquired in somatic tissues can enter the germline and be transmitted through sexual reproduction, we compared eccDNA profiles of the graft-derived sexual progenies cp-TTT2 and rcp-TTT1 with those of the self-grafted controls rgp-TTT and rgp-CCC. EccDNA detected in sexual progeny but absent from rgp-TTT was first identified, and its presence in rgp-CCC was subsequently examined to infer its putative donor-genome origin (Fig. 3a). This comparative framework enabled the lineage-resolved inference of heterologous, somatically acquired eccDNAs entering the reproductive lineage.

**Fig. 3.**
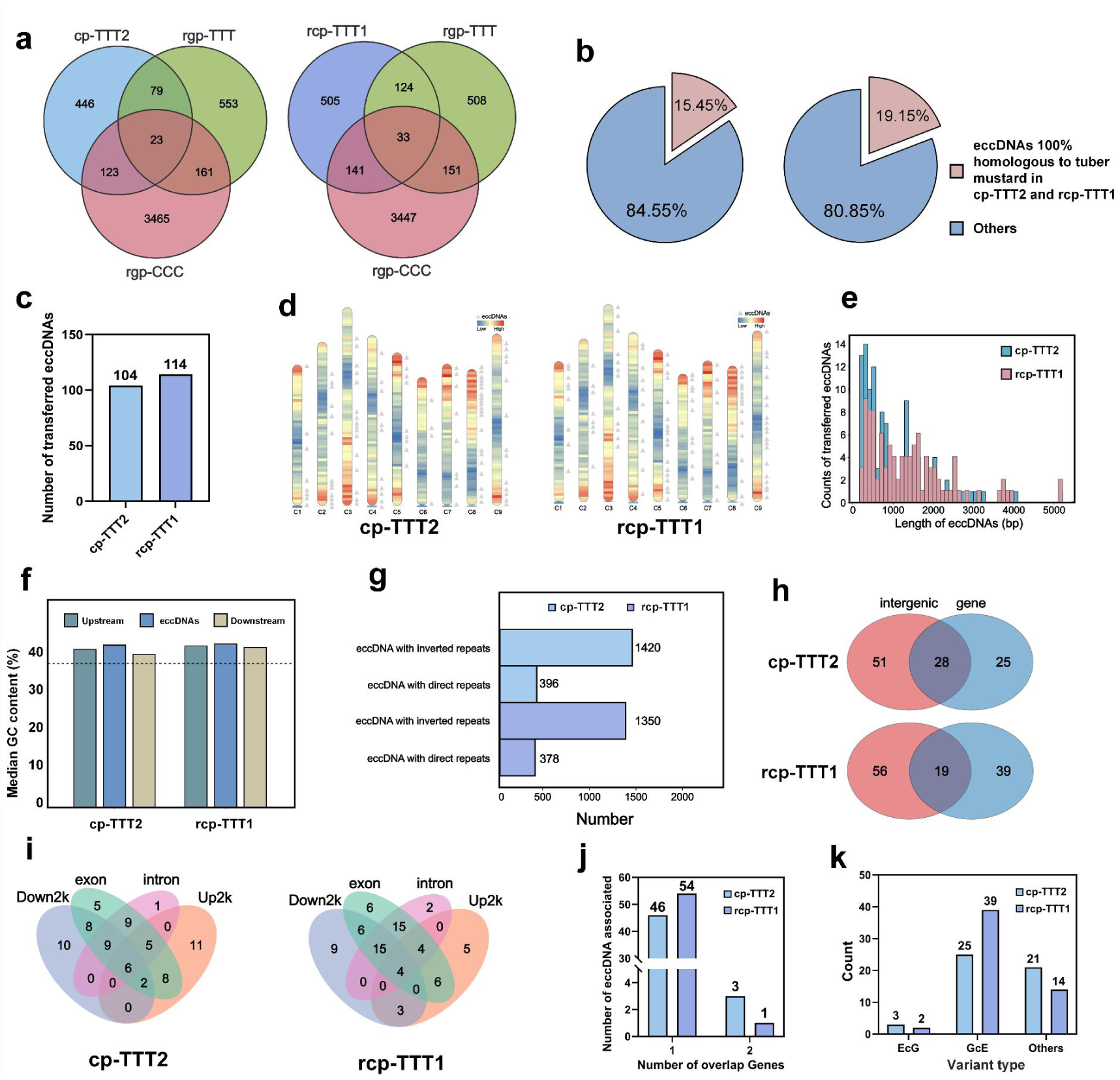
Identification and sequence characterization of transmitted eccDNAs in the sexual progeny of Brassica chimeras. **a,** Venn diagrams displaying the overlaps of eccDNA loci mapped to the *B. oleracea* reference genome among cp-TTT2, rcp-TTT1, rgp-TTT, and rgp-CCC. **b,** Among eccDNAs mapped to the *B. oleracea* reference genome, the fraction showing 100% sequence identity to *B. juncea* versus other sequences. **c,** Number of transmitted *B. oleracea*–specific eccDNA detected in cp-TTT2 and rcp-TTT1. **d,** Chromosomal distribution of transmitted *B. oleracea*-specific eccDNAs (n=104 in cp-TTT2; n=114 in rcp-TTT1). Red: high gene density; blue: low gene density; triangles: locations of eccDNA on the *B. oleracea* genome. **e,** Length distribution of transmitted eccDNAs in cp-TTT2 and rcp-TTT1. **f,** Median GC content of transmitted eccDNAs compared with 150-bp flanking regions upstream and downstream of their source loci; the genome-wide mean GC content of *B. oleracea* is shown as a dashed line. **g,** Number of transmitted eccDNA containing direct or inverted repeats. **h,** Number of transmitted eccDNA overlapping genic or intergenic regions. **i,** Number of transmitted eccDNA overlapping with exons, introns, and ±2-kb genic flanks (up2kb and down2kb). **j,** Count of transmitted eccDNAs associated with one gene or spanning two genes. **k,** Comparison of EcG and GcE variant types in cp-TTT2 and rcp-TTT1 (as defined in the main text).

In cp-TTT2, 569 eccDNAs were uniquely detected relative to rgp-TTT, of which 123 could be assigned to *B. oleracea* based on their presence in rgp-CCC (Fig. 3a; Extended Data Fig. 2), indicating stable retention of graft-acquired *B. oleracea*–derived eccDNA in the sexual progeny. A comparable pattern was observed in rcp-TTT1, which harbored 646 unique eccDNAs, including 141 shared with rgp-CCC (Fig. 3a; Extended Data Fig. 2). After excluding eccDNA showing 100% sequence identity to *B. juncea*, 104 and 114 *B. oleracea*–specific eccDNA molecules were retained in cp-TTT2 and rcp-TTT1, respectively (Fig. 3b–c; Extended Data Table 4). These *B. oleracea*–specific transmitted eccDNAs represented 84.55% and 80.85% of the *B. oleracea*–mapped eccDNA sets in the two sexual progenies, providing strong evidence for graft-mediated horizontal transfer followed by stable inheritance through the germline.

Genome-wide mapping revealed that transferred eccDNA was broadly distributed across the *B. oleracea* genome (Fig. 3d). EccDNA occurrence showed a pronounced bias toward gene-dense chromosomal regions, particularly near one or both chromosomal ends (Fig. 3d). Overall, the distribution suggests that eccDNA excision is not random but constrained by local genomic context. Notably, chromosomes C8 and C9 displayed markedly elevated eccDNA frequencies in regions of high gene density (Fig. 3d), consistent with observations in animal systems showing preferential eccDNA formation in gene-rich domains.

The total number of transferred eccDNA was slightly higher in rcp-TTT1 than in cp-TTT2, potentially reflecting additional genome restructuring during tissue regeneration prior to sexual reproduction ^43^. The size distribution of transferred eccDNA ranged from 139 to 4,136 bp in cp-TTT2 and from 139 to 5,212 bp in rcp-TTT1, with a pronounced enrichment between 200 and 800 bp in both progeny lines (Fig. 3e; Extended Data Table 5). This size profile closely mirrors those reported in human cells ^44^, *Arabidopsis* ^44^, and rice ^20,21^, suggesting conserved constraints on eccDNA architecture across the kingdoms. In agreement with previous plant studies, eccDNA and their 150-bp flanking regions exhibited elevated GC content relative to the genomic background, with the highest enrichment observed within the eccDNA sequences themselves (Fig. 3f; Extended Data Table 6) ^21,26^.

To identify sequence features associated with the biogenesis and persistence of acquired eccDNA, we systematically analyzed eccDNA termini together with ±150-bp flanking sequences. BLAST analyses revealed a pronounced enrichment of both inverted and direct repeat motifs at eccDNA termini, a feature previously implicated in eccDNA excision and circularization in both animal and plant systems ^21,22^. In cp-TTT2, 1,420 inverted repeat pairs were identified among 100 transferred eccDNAs, whereas in rcp-TTT1, 1,350 inverted repeat pairs were detected in 113 of the 114 transferred eccDNA molecules (Fig. 3g; Extended Data Table 7). Similarly, 396 direct repeat pairs were identified in 102 transferred eccDNAs in cp-TTT2, and 378 direct repeat pairs were detected in 110 of 114 transferred eccDNAs in rcp-TTT1 (Fig. 3g; Extended Data Table 8). Notably, the vast majority of eccDNA contained multiple inverted and direct repeat pairs, with repeat lengths tightly constrained to 6–8 bp (Extended Data Tables 7 and 8). The strong convergence of these genetic features across independent graft lineages indicates conserved eccDNA architecture, consistent with a role in excision, circularization, and long-term maintenance ^21^.

To characterize the genomic distribution of eccDNA inherited in sexual progeny, we mapped their loci to specific genomic compartments. In cp-TTT2, 53 eccDNAs overlapped with annotated gene regions and 79 with intergenic regions; in rcp-TTT1, 58 were gene-associated and 75 intergenic (Fig. 3h; Extended Data Table 9). These proportions are consistent with reports in rice, where eccDNA is enriched in intergenic regions and underrepresented within gene bodies ^21^. Notably, 28 eccDNAs in cp-TTT2 and 19 in rcp-TTT1 spanned both genic and intergenic regions, indicating that acquired eccDNA is not confined to discrete genomic compartments. Among gene-associated eccDNA, 97–98% overlapped exons, with only a small fraction mapped to introns (Fig. 3i; Extended Data Table 10). For intergenic eccDNA, substantial proportions overlapped putative upstream (−2 kb) or downstream (+2 kb) regulatory regions of annotated genes (Fig. 3i; Extended Data Table 10), suggesting potential regulatory roles, as proposed in previous functional analyses of plant eccDNA ^21,44^.

Applying a stringent criterion requiring ≥30% overlap with annotated gene regions, we identified 46 and 54 gene-associated eccDNAs in cp-TTT2 and rcp-TTT1, respectively (Fig. 3j; Extended Data Table 11). In further analyses, we categorized gene containing whole eccDNA as ‘GcE’ and eccDNA containing whole gene as ‘EcG’. Most eccDNA corresponded to a single gene, reflecting their generally short length, and were classified as GcEs, whereas only a small fraction of eccDNA encompassed entire genes (Fig. 3k; Extended Data Table 12). Notably, a limited number of EcGs encompassed intact protein-coding genes. Three in cp-TTT2 (*Boc09g04685, Boc03g03075* and *Boc09g00867*) and two complete genes were detected in rcp-TTT1 (*Boc09g02204* and *Boc04g02259*). Functional annotation based on homology to genes in *Arabidopsis thaliana* and other Brassicaceae species indicates that these genes encode conserved proteins involved in nuclear regulation, RNA metabolism, stress responses, and core metabolic processes (Extended Data Table 3).

Collectively, these results demonstrate that eccDNAs acquired in somatic tissues can traverse the somatic–germline boundary and be stably inherited by sexual progeny. The observed non-random genomic distribution, conserved size constraints, and enrichment of inverted and direct repeat motifs indicate that eccDNA detected in sexual progeny possesses intrinsic features compatible with germline persistence, echoing models of eccDNA selection and maintenance proposed in animal and plant systems ^20–22^. These findings support a model in which eccDNA functions as a mobile carrier of genetic information, providing a route for graft-associated somatic genetic exchange to contribute to heritable variation. We therefore next examined whether the transmitted eccDNA harbors conserved motifs associated with autonomous replication and transcriptional activity.

### Heritable eccDNA carries intrinsic signatures of autonomous replication

The stable transmission of eccDNA in sexual progeny prompted us to examine whether graft-acquired eccDNA retains intrinsic signatures associated with autonomous replication. We systematically searched for the canonical 11-bp ACS motif in inherited eccDNA of periclinal chimera progeny. 13 conserved origin-like motifs were identified across 9 eccDNAs in cp-TTT2, whereas 18 motifs were detected across 12 eccDNAs in rcp-TTT1 (Fig. 4a, b; Extended Data Table 13). In rcp-TTT1, one eccDNA (C8:5975352–5976845; 1,493 bp) contained three ACS motifs. Four other eccDNA molecules harbored two motifs near termini, and the remaining eccDNA carried a single motif. A similar pattern was observed in cp-TTT2, where one eccDNA (C2:27966257–27967600; 1,343 bp) contained three motifs, three eccDNA molecules had two motifs, and six molecules carried one (Extended Data Table 13). The non-random enrichment of ACS motifs, particularly their recurrent localization near eccDNA terminal ends, or within larger circular molecules, is consistent with non-random retention of replication-associated sequence features.

**Fig. 4.**
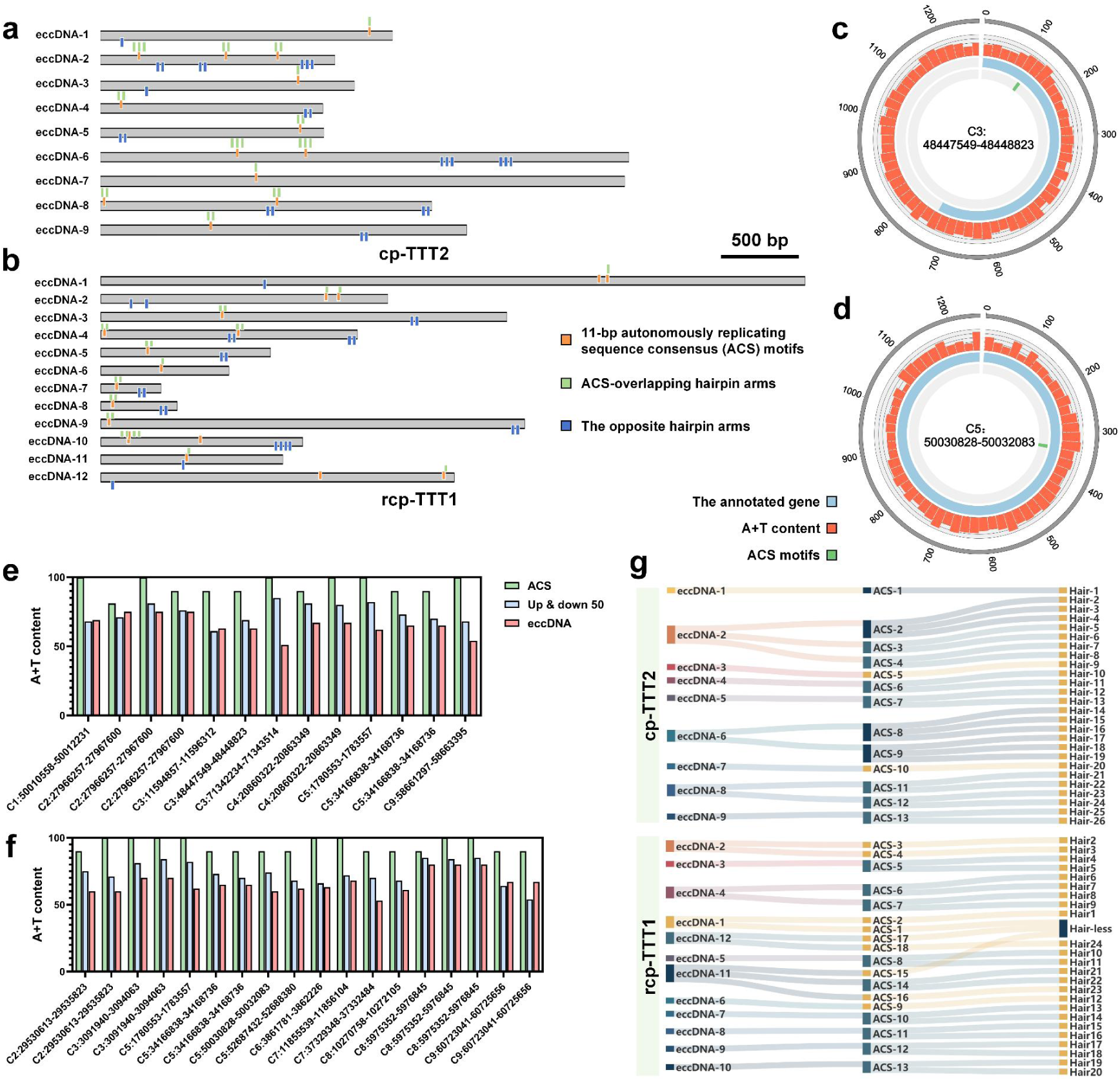
Analysis of autonomously replicating sequence consensus (ACS) and hairpin structures in transmitted eccDNAs from Brassica chimera progeny. **a,b,** ACS-like 11-bp motifs and predicted hairpin arms in transmitted eccDNAs detected in cp-TTT2 (a; n=9 eccDNAs) and rcp-TTT1 (b; n=12 eccDNAs). Orange markers denote ACS-like 11-bp motifs; green markers indicate ACS-overlapping hairpin arms; blue markers indicate the opposing hairpin arms. **c,d,** Circular maps of representative transmitted eccDNAs from cp-TTT2 (c; C3: 48,447,549–48,448,823) and rcp-TTT1 (d; C5: 50,030,828–50,032,083). Outer circles indicate genomic coordinates; red tracks show A+T content (50-bp sliding windows); blue blocks indicate annotated genes (*Boc03g03414* in c; *Boc05g03674* in d); green ticks indicate ACS-like motif positions. **e,f,** A+T content of ACS-like motifs (11 bp), their immediate 50-bp upstream and downstream flanks, and full-length eccDNAs in cp-TTT2 (e) and rcp-TTT1 (f). **g,** Mapping between ACS-like motifs and associated hairpin elements in transmitted eccDNAs, comprising 13 motifs and 26 hairpin elements in cp-TTT2 and 18 motifs and 24 hairpin elements in rcp-TTT1.

Notably, several ACS motifs were embedded within protein-coding regions. In cp-TTT2, an eccDNA derived from chromosome 3 (C3:48447549–48448823) contained an ACS motif within the gene *Boc03g03414* (Fig. 4c; Extended Data Table 14). In rcp-TTT1, five ACS motifs were located within genes (*Boc05g03674*, *Boc05g03996*, *Boc07g01715*, *Boc07g03746* and *Boc08g01595*; Fig. 4d; Extended Data Fig. 3a–d; Extended Data Table 14). This intra-gene positioning mirrors that of the eccDNA replicon described in *A. palmeri*, in which replication initiates within a NAC-domain transcription factor ^45^. These observations raise the possibility that some plant eccDNAs harbour origin-like motifs within gene bodies, although whether this positioning influences replication competence or persistence remains to be determined. Additional ACS motifs were also identified in intergenic regions (Extended Data Table 14).

Consistent with the AT-rich nature of the ACS consensus, ACS motifs displayed pronounced local A+T enrichment. In rcp-TTT1, A+T content across the 18 ACS motifs ranged from 90.91% to 100%, whereas in cp-TTT2, 12 of 13 motifs fell within the same range, with a single exception at 81.82% (Fig. 4e, f; Extended Data Table 15). By contrast, A+T content declined sharply within the 50 bp region flanking the ACS motifs, and further across entire eccDNA molecules (Fig. 4e, f; Extended Data Table 15). This local enrichment supports the interpretation that these ACS-like motifs reside in sequence contexts compatible with origin-like activity on transmitted eccDNAs.

Beyond primary sequence motifs, secondary DNA structure is increasingly recognized as a critical facilitator of circular DNA replication. Inverted repeats and hairpin-forming sequences can provide intrinsic priming sites for rolling-circle amplification (RCA), a mechanism widely used by viral replicons, plasmids, and organellar genomes ^29,35–37^. In plant organelles, origins of replication often coincide with inverted repeat regions, and origin-independent replication can be initiated via short repeats or R-loop–associated priming structures ^46–48^. These diverse systems underscore the importance of secondary DNA architecture in sustaining circular genome replication.

We therefore examined whether heritable eccDNA exhibits analogous structural features. In cp-TTT2, all 104 eccDNAs transmitted to sexual progeny contained predicted hairpin structures or inverted repeats, yielding a total of 7,111 paired secondary-structure elements (Extended Data Table 16). Integrating motif and structure annotations revealed strong spatial coupling between ACS-like motifs and secondary-structure features. Strikingly, all 13 ACS regions in cp-TTT2 overlapped with hairpin structures or inverted repeats, and 26 hairpin arms directly spanned the 11-bp core sequence (Fig. 4b and 4g; Extended Data Table 17).

A comparable pattern was observed in rcp-TTT1. Among 114 heritable eccDNAs, 7,086 paired hairpin or inverted-repeat structures were identified (Extended Data Table 16), and and 17 of 18 ACS-containing regions overlapped with these secondary structures (Extended Data Table 17). In total, 24 hairpin arms coincided with ACS cores, and in one eccDNA, two ACS regions were entirely encompassed within hairpin structures (Fig. 4a and 4g; Extended Data Tables 16 and 17). The consistent coupling between origin-like motifs and secondary DNA structures across independent progenies suggests that ACS-like motifs frequently reside within hairpin-forming contexts, a configuration that could facilitate priming and contribute to eccDNA persistence after germline transmission.

### Reproducible yet divergent eccDNA inheritance among sexual progenies

To evaluate the reproducibility and lineage-dependent divergence of somatically acquired eccDNA inheritance, we performed comparative profiling of transferred eccDNAs across biological replicates of cp-TTT2 and rcp-TTT1. In cp-TTT2, 98 and 73 eccDNA molecules were detected in the two biological replicates, whereas 112 and 82 were identified in the two rcp-TTT1 replicates (Fig. 5a; Extended Data Table 18). Across all samples, eccDNA and their 150-bp flanking regions exhibited GC contents consistently higher than the genomic average, with eccDNA showing the strongest enrichment (Fig. 5b; Extended Data Table 18), in agreement with previous observations ^20–22^. Length distribution analysis revealed that most eccDNA ranged from 200 to 800 bp, and similar size profiles were observed between biological replicates within each progeny line (Extended Data Fig. 4a; Extended Data Table 18), underscoring the robustness of eccDNA detection and inheritance patterns.

**Fig. 5.**
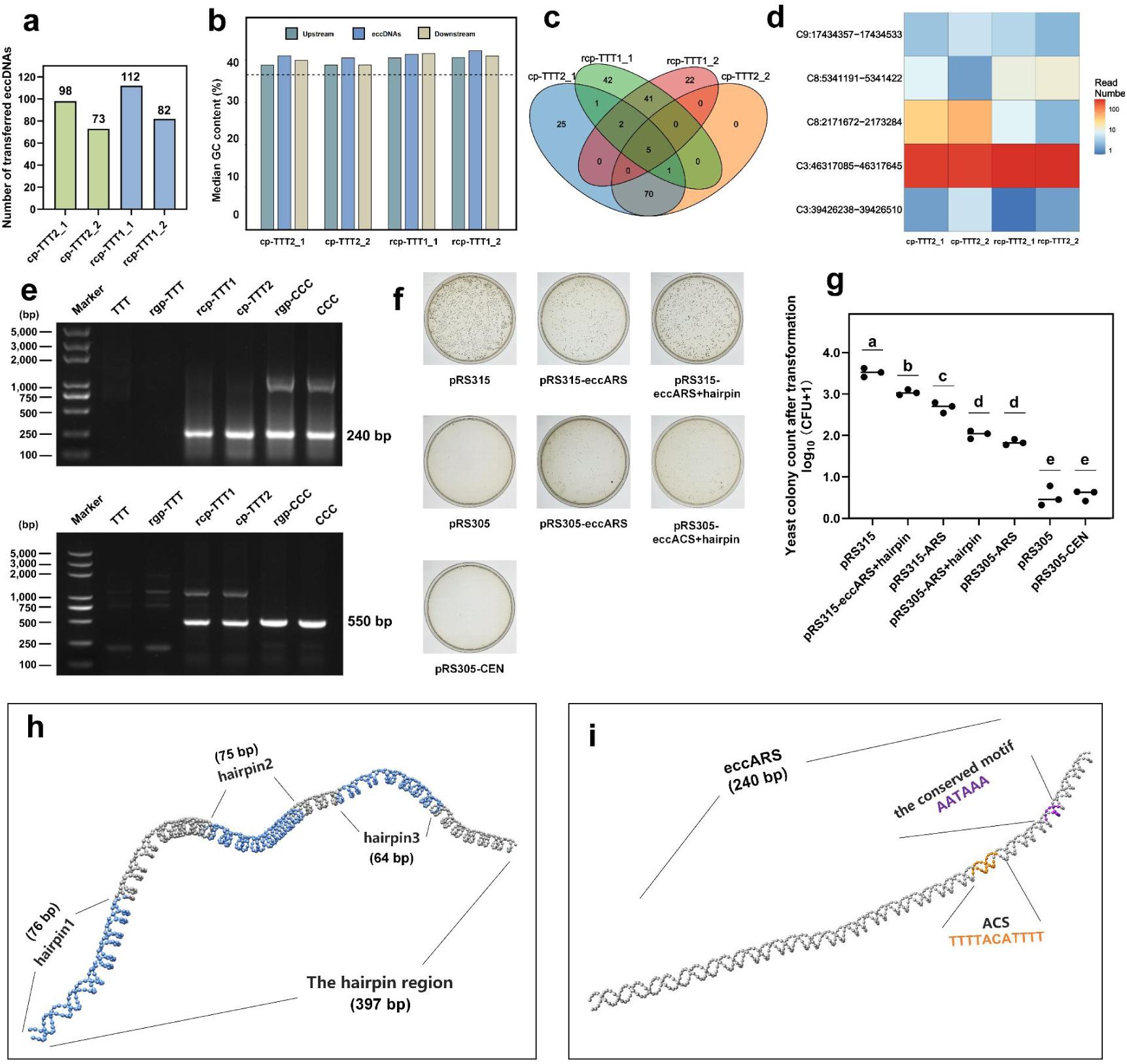
Consistency of transmitted eccDNAs and replication competence in yeast. **a,** Numbers of transmitted *B. oleracea*–specific eccDNAs detected in biological replicates of cp-TTT2 (cp-TTT2_1, cp-TTT2_2) and rcp-TTT1 (rcp-TTT1_1, rcp-TTT1_2). **b,** Median GC content of transmitted eccDNAs compared with 150-bp upstream and downstream flanks at their source loci; the genome-wide mean GC content of *B. oleracea* is shown as a dashed line. **c,** Overlap of transmitted eccDNA sets between biological replicates of cp-TTT2 and rcp-TTT1. **d,** Supporting read counts for five eccDNAs detected in both progeny lines and both replicates. **e,** PCR validation of two representative transmitted eccDNAs (C3:46,317,085–46,317,645 and C5:42,317,656–42,319,060) across the indicated samples (TTT, rgp-TTT, rcp-TTT1, cp-TTT2, rgp-CCC and CCC). **f,** Yeast assay testing whether eccDNA-derived sequences support maintenance of ARS-deficient plasmids. Shown are representative plates for the indicated constructs with or without eccARS and an adjacent hairpin-containing fragment in pRS315ΔARS and pRS305 backbones. **g,** Colony-forming units (CFU) after transformation with the indicated constructs; dots denote independent transformations. Different letters indicate significant differences (P < 0.05, Tukey’s HSD). **h,** Predicted secondary-structure features within a 397-bp hairpin-rich region of the eccDNA replicon, highlighting three hairpin elements (76 bp, 75 bp and 64 bp). **i,** Predicted structural features within a 240-bp eccARS-containing region, marking the ACS-like motif (orange) and an AATAAA-containing element (purple).

Within each progeny line, inheritance of somatically acquired eccDNA was highly reproducible. 76 were shared between the two cp-TTT2 replicates, and 48 were shared between the two rcp-TTT1 replicates (Fig. 5c). Notably, five eccDNA molecules were consistently detected in both replicates of cp-TTT2 and rcp-TTT1 (Fig. 5c). Although these eccDNAs were shared across independent lineages, their accumulation levels differed markedly (Fig. 5d; Extended Data Table 19). Quantitative analysis of read abundance revealed that one eccDNA spanning C3:46317085–46317645 accumulated to substantially higher levels across all four samples relative to the other shared eccDNAs (Fig. 5d). Within this shared set, several eccDNAs showed reproducible differences in abundance between cp-TTT2 and rcp-TTT1 replicates. For example, eccDNA corresponding to C3:39426238–39426510 and C8:2171672–2173284 were consistently more abundant in both cp-TTT2 replicates than in rcp-TTT1. Conversely, eccDNA located at C3:46317085–46317645 and C8:5341191–5341422 accumulated to lower levels in cp-TTT2 than in rcp-TTT1, whereas C9:17434357–17434533 displayed intermediate abundance between the two progeny lines (Fig. 5d). These results indicate that eccDNA inheritance is not only consistent but also quantitatively modulated in a lineage-dependent manner, suggesting selective maintenance rather than passive carryover.

To independently validate eccDNA transmission, we performed junction-specific PCR for two representative eccDNA molecules selected at random. One eccDNA located at C3:46317085–46317645 was undetectable in the *B. juncea* parental control (TTT) and its self-grafted sexual progeny (rgp-TTT) yet yielded clear junction-specific amplicons in both rcp-TTT1 and cp-TTT2, as well as in the *B. oleracea* parental control (CCC) and its self-grafted progeny (rgp-CCC) (Fig. 5e; Extended Data Table 20; Supplemental Information 1). This eccDNA encompassed a gene exhibiting ∼70% sequence identity to *A. thaliana ERF4* and >88% identity to the *B. oleracea ERF4* reference (Extended Data Fig. 4b). Conserved motif analysis confirmed the presence of canonical ERF domains (Extended Data Fig. 4c), indicating functional conservation. Given links between ERF4-family regulators and drought-related traits ^49,50^, this eccDNA provides a candidate locus for connecting eccDNA retention with the drought-tolerance phenotype observed in chimera progenies.

A second shared eccDNA, spanning C5:42317656–42319060, exhibited a similar inheritance pattern, absent from TTT and rgp-TTT but consistently detected in both graft-derived progeny lines as well as in CCC and rgp-CCC controls (Fig. 5e; Extended Data Table 20). Sanger sequencing confirmed precise circular junctions but also revealed an additional ∼240 bp insertion relative to the genomic mapping interval (Supplementary information 2). This additional fragment originated from a distinct locus on the same chromosome and harbored a conserved 11-bp ACS-like motif, suggesting that recombination or fusion events may generate composite eccDNA structures that carry replication-associated modules. Consistent with this interpretation, accumulating evidence indicates that eccDNA is highly dynamic, capable of recombination and fusion, giving rise to structurally complex circular DNAs through homology-mediated end joining and repeat-driven rearrangements in both normal and pathological contexts ^22,51,52^.

To test whether the identified ACS-like-containing sequence contributes to eccDNA replication and maintenance, we employed two complementary yeast plasmid systems: the replicating pRS315 plasmid, which contains endogenous centromere (CEN) and ARS elements, and a non-replicating plasmid backbone lacking autonomous replication sequences (pRS305). To specifically assess the replication activity of the eccDNA-derived ARS-like sequence (C5:42317656–42319060), the endogenous ARS element was deleted from pRS315 to generate the ARS-deficient pRS315ΔARS vector (Fig. 5f and 5g; Extended Data Table 20). As expected, the intact pRS315 plasmid carrying the native ARS and CEN6 elements produced dense colonies following transformation, reflecting robust autonomous replication. Insertion of the 240 bp eccDNA-derived fragment into pRS315ΔARS supported colony formation above background, indicating that this element can substitute for a missing origin in yeast (Fig. 5f and 5g). Notably, inclusion of an adjacent 397 bp region composed of three tandem repeats (76 bp, 75 bp and 64 bp) enriched in predicted hairpin-forming structures further increased colony formation relative to the 240 bp fragment alone (Fig. 5f and 5g), suggesting that local secondary structures enhance replication efficiency. Nevertheless, replication driven by these eccDNA-derived elements remained substantially weaker than that conferred by the endogenous yeast ARS.

To further dissect the contribution of the centromeric element to eccDNA maintenance, we performed parallel assays using the pRS305 plasmid, which lacks autonomous replication elements (Fig. 5f and 5g; Extended Data Table 20). Introduction of the CEN6 sequence alone into pRS305 failed to support colony formation, indicating that the centromere element is insufficient to drive replication in the absence of a functional origin. The empty pRS305 vector similarly yielded no colonies and served as a negative control. In contrast, insertion of the 240 bp fragment enabled limited colony formation, although at a lower frequency than observed in the pRS315ΔARS background, consistent with replication activity that is independent of vector context but influenced by episomal content. As observed in the pRS315 system, addition of the adjacent hairpin-containing region modestly enhanced colony formation compared with the 240 bp fragment alone yet remained less effective than the corresponding construct in the pRS315 background (Fig. 5f and 5g). Collectively, these results demonstrate that the ARS-like sequence identified from eccDNA confers autonomous replication capacity in yeast, that adjacent hairpin-forming regions enhance replication efficiency, and that the centromeric element contributes to episomal stability but cannot substitute for a functional origin of replication. These assays therefore provide functional support for cis features that could facilitate eccDNA persistence after transmission.

Plasmid retention was further validated by serial passaging on selective medium followed by colony PCR, which detected eccDNA-derived origin-like sequence exclusively in yeast transformed with eccDNA-derived constructs and not in vector-only controls (Extended Data Fig. 5). Consistent with these functional assays, DNA curvature modeling of the 397 bp hairpin-containing region revealed pronounced and sustained DNA bending across hairpin1–hairpin3, accompanied by localized sharp bends indicative of reduced helical stability (Fig. 5h), features previously associated with origin-proximal regions ^45^. A comparable bending profile was observed within the 240 bp ARS-containing fragment encompassing the conserved 11-bp motif and a predicted DNA unwinding element (Fig. 5i).

Together, these findings demonstrate that eccDNA acquired through somatic grafting can be stably and reproducibly transferred to sexual progeny, while exhibiting lineage-specific accumulation dynamics. The identification and functional validation of replication-associated cis-elements provide mechanistic support for selective eccDNA maintenance and support eccDNA retention as a contributor to heritable phenotypic diversification following shoot apical meristem grafting.

### Heritable phenotypic diversification associated with eccDNA transmission

Following the transmission of somatically acquired eccDNA into the reproductive lineage, both sexual progeny lines exhibited pronounced, heritable phenotypic alterations relative to the self-grafted parental control. All observed variations demonstrated a degree of directionality and consistency across plants. The most conspicuous changes were observed in leaf morphology and size (Fig. 6a, Extended Data Fig. 6). Compared with rgp-TTT, leaves of rcp-TTT1 and cp-TTT2 displayed significantly reduced length-to-width ratios (by 32.20% and 31.69%, respectively), accompanied by a concomitant reduction in total leaf area (by 11.05% and 10.23%, respectively) (Fig. 6b-e). These quantitative changes were further associated with visibly shallower marginal serration, yielding leaf architectures that more closely resembled those of rgp-CCC than those of the *B. juncea* parental type (Fig. 6a, Extended Data Fig. 6). Importantly, these morphological traits were maintained with high fidelity over more than six successive generations, demonstrating long-term stability of graft-associated phenotypic variation.

**Fig. 6.**
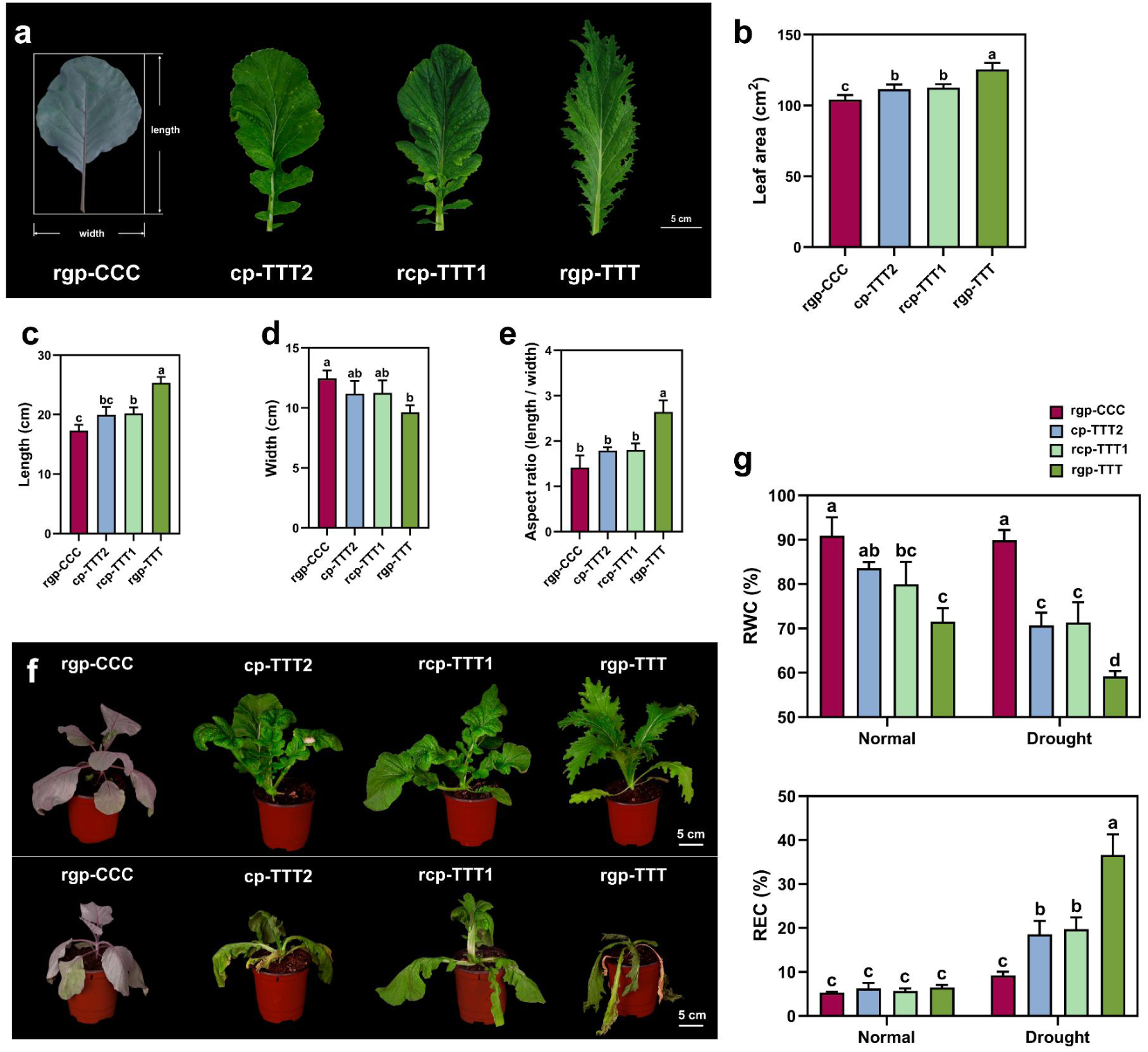
Graft-induced variation in leaf traits and drought response in chimera-derived sexual progeny. **a,** Representative leaves showing the measurement scheme (length and width) and leaf morphology of rgp-CCC, cp-TTT2, rcp-TTT1 and rgp-TTT (left to right). Scale bar, 5 cm. **b–e,** Quantification of leaf area (b), leaf length (c), leaf width (d) and aspect ratio (length/width; e) for the indicated genotypes. **f,** Representative whole-plant images of rgp-CCC, cp-TTT2, rcp-TTT1 and rgp-TTT grown under control conditions (top) or drought treatment (bottom). Scale bar, 5 cm. **g,** Relative water content (RWC) and relative electrical conductivity (REC) measured under control conditions and after 7 d drought (water withholding) for the indicated genotypes.

Given the drought tolerance of *B. oleracea*, we next assessed whether eccDNA-containing progenies carrying somatically acquired eccDNA exhibited altered responses to water deficit. Under drought conditions, rgp-TTT plants showed severe wilting, whereas rgp-CCC, rcp-TTT1 and cp-TTT2 retained leaf turgor to varying degrees (Fig. 6f). Histological analysis further revealed increased leaf thickness in rgp-CCC, rcp-TTT1 and cp-TTT2 relative to rgp-TTT after 7 days of drought treatment (Extended Data Fig. 7), suggesting enhanced tissue robustness. Consistent with these anatomical differences, physiological measurements placed rcp-TTT1 and cp-TTT2 between rgp-TTT and rgp-CCC with respect to relative water content and electrolyte leakage (Fig. 6g), indicating improved cellular water retention and membrane stability under stress.

These results demonstrate that sexual progeny derived from periclinal chimeras inherit stably intermediate morphological and physiological traits between the two parental species. The convergence of leaf architecture and drought-response phenotypes in cp-TTT2 and rcp-TTT1, despite their distinct developmental origins, is consistent with a shared, heritable change acquired in the chimera context. In parallel with eccDNA retention in these progenies, the phenotypic stability provides a framework to explore links between graft-associated genetic exchange and durable trait variation in *B. juncea*.

### Transcriptionally active eccDNA reshapes gene expression networks

To determine whether the inheritance of heterologous somatically acquired eccDNA is associated with coordinated changes in gene expression, RNA sequencing was performed on sexual progenies of self-grafted chimera controls (rgp-TTT) and the two chimeric-derived progeny lines (cp-TTT2 and rcp-TTT1) at the four-leaf stage. Both progeny lines exhibited extensive transcriptional reprogramming relative to rgp-TTT (Fig. 7a). cp-TTT2 displayed 5,247 differentially expressed genes (DEGs), comprising 2,976 upregulated and 2,271 downregulated genes, whereas in rcp-TTT1, 7,186 DEGs were identified, including 4,191 upregulated and 2,995 downregulated genes (Fig. 7a).

**Fig. 7.**
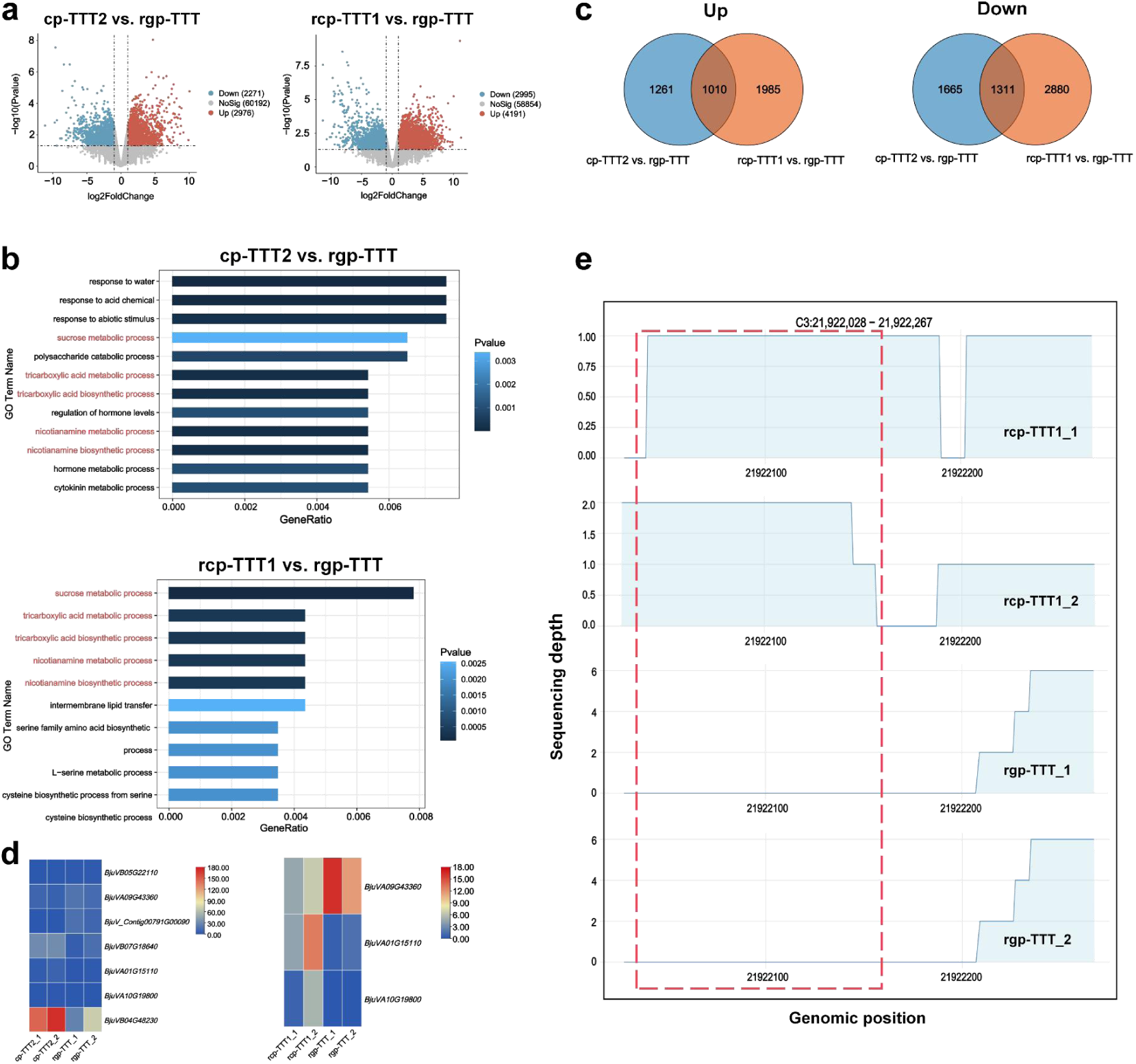
Transcriptional changes in eccDNA-containing Brassica progeny. **a,** Volcano plots of differential gene expression for cp-TTT2 versus rgp-TTT and rcp-TTT1 versus rgp-TTT. Differential expression was assessed using edgeR; genes with |fold change| ≥ 1.5 and FDR < 0.05 (Benjamini–Hochberg) are highlighted (red, upregulated; blue, downregulated). **b,** Gene Ontology (GO) biological process enrichment analysis for DEGs in cp-TTT2 versus rgp-TTT (top) and rcp-TTT1 versus rgp-TTT (bottom). Enrichment *P* values were calculated using a hypergeometric test and adjusted using Benjamini–Hochberg; GO terms enriched in both comparisons are highlighted in red. **c,** Overlap of upregulated (left) and downregulated DEGs identified in cp-TTT2 versus rgp-TTT and rcp-TTT1 versus rgp-TTT. **d,** Heatmap of selected DEGs related to leaf development and drought response across cp-TTT2, rcp-TTT1 and rgp-TTT (two biological replicates per line). Values are row-scaled z-scores. **e,** Coverage tracks showing RNA-seq signal at an eccDNA-associated locus (C3: 43,598,797–43,600,805) in rcp-TTT1_1/2 and rgp-TTT replicates.

A substantial core response was shared between cp-TTT2 and rcp-TTT1 relative to rgp-TTT at both the gene and pathway levels. A total of 1,311 genes were commonly upregulated and 1,010 genes commonly downregulated in both cp-TTT2 and rcp-TTT1 relative to rgp-TTT (Fig. 7c). These shared upregulated genes were significantly enriched in sucrose metabolism, nicotianamine metabolism, and the tricarboxylic acid (TCA) cycle (Fig. 7b), forming a metabolic core that supports carbon allocation, metal homeostasis, and energy production. Enhanced sucrose metabolism contributes to osmotic adjustment and carbon supply ^53^, while nicotianamine-mediated metal chelation can mitigate oxidative stress ^54^, and activation of the TCA cycle sustains ATP production and redox balance ^55^. Together, these pathways are consistent with the coordinated changes in growth, leaf architecture, and stress resilience observed in both progeny lines.

To link transcriptional changes to altered leaf morphology, we examined the expression of key regulators of organ development and polarity. In cp-TTT2, *BjuVB05G22110* (*KANADI*) expression was reduced to 37.03% of the control (Fig. 7d), consistent with the role of *KAN* genes in establishing abaxial identity and leaf polarity ^56^. In rcp-TTT1, expression of *BjuVA09G43360* (*CYCD3;3*), a cytokinin-responsive regulator of cell proliferation in lateral organs ^56^, was reduced to 38.64% of control levels (Fig. 7d). Notably, *CYCD3;3* expression was similarly suppressed in cp-TTT2, along with *BjuV_Contig00791G00090* (*CYCD3;3*), indicating convergent transcriptional modulation of leaf-shape–associated genes across progenies derived from distinct chimeric tissues. These transcriptional expression patterns closely corresponded to the altered leaf morphologies observed in rcp-TTT1 and cp-TTT2 (Fig. 6a–e).

Genes involved in drought responses were also consistently activated in eccDNA-containing progenies. In cp-TTT2, *BjuVB07G18640*, encoding an MYB96-like transcription factor, was upregulated 2.89-fold relative to rgp-TTT (Fig. 7d). MYB96 is known to promote cuticular wax biosynthesis and enhance drought tolerance ^57^. Similarly, *BjuVB04G48230* (*NAC019*) showed a 2.92-fold increase in cp-TTT2, consistent with its established role in stress-responsive transcriptional regulation ^58^. In rcp-TTT1, *BjuVA01G15110* (*AIRP1*), encoding a C3H2C3-type RING E3 ubiquitin ligase involved in ABA-dependent drought responses, was strongly induced (7.94-fold), with a slightly weaker induction also detected in cp-TTT2 (4.46-fold; Fig. 7d) ^59^. In addition, *BjuVA10G19800* (*CBL5*), whose overexpression confers drought tolerance ^60^, was markedly upregulated in both rcp-TTT1 (32.26-fold) and cp-TTT2 (8.02-fold). The consistent activation of these drought-responsive genes suggests enhanced stress resilience in both progeny lines relative to the control.

Differences between the two progeny lines were also observed. DEGs uniquely upregulated in cp-TTT2 were enriched for cytokinin and hormone-related processes, abiotic stress responses, and polysaccharide catabolism (Fig. 7c) ^61,62^, whereas rcp-TTT1-specific DEGs were enriched for serine/cysteine biosynthesis, amino acid metabolism, and intermembrane lipid transfer (Fig. 7b), consistent with redox buffering and membrane remodeling ^63^. The larger number of DEGs observed in rcp-TTT1 may reflect its distinct developmental origin, as rcp-TTT1 was derived from the regenerants of the L1 cell layer of the chimera, whereas cp-TTT2 directly derived from the L2 cell layer of the chimeric organism.

To directly assess whether transferred eccDNAs are transcriptionally active, we interrogated transcriptome datasets against eccDNA sequences identified in the progenies. Transcripts uniquely mapping to eccDNA were detected in cp-TTT2 and rcp-TTT1 but were absent from rgp-TTT controls (Fig. 7e and Extended Data Table 22). In cp-TTT2, 18 of 104 transferred eccDNA molecules produced eccDNA-specific transcripts, whereas 12 of 114 in rcp-TTT1 showed evidence of transcriptional activity (Fig. 7e; Extended Data Fig. 8 and 9; Extended Data Table 22). These results indicate that a subset of transferred eccDNA is transcriptionally competent in the sexual progeny and may directly contribute to gene-expression landscapes.

Collectively, these results link eccDNA retention in chimera-derived progeny with widespread transcriptional changes. The convergence of widespread transcriptional reprogramming, modulation of key developmental and stress-responsive regulators, and direct transcription from eccDNA is consistent with a contribution of inherited eccDNA to the altered transcriptional landscape in sexual progeny.

## Discussion

During evolution, plants must continuously generate and maintain genetic variation to enhance their adaptability. Extrachromosomal circular DNA (eccDNA) is increasingly recognized as a dynamic source of genomic variation, capable of being induced by environmental cues and persisting as stable extrachromosomal elements ^17,20,64^. Throughout the plant life cycle, eccDNAs can arise endogenously and, notably, can also be transferred and exchanged between individuals through grafting and potentially other interplant pathways. We previously showed that eccDNA can move horizontally between heterologous tissues in grafted plants and remain maintained in recipient somatic cells ^26^. Here, using lineage-resolved periclinal chimeras, we addressed whether graft-acquired eccDNA can cross the soma–germline boundary. We showed that exogenous eccDNA can access germline lineages and be passed onto sexual progeny across multiple generations. Inherited eccDNA is enriched for cis-encoded sequences and structural features linked to episomal persistence and displayed replication competence in a heterologous assay. Moreover, a subset of inherited eccDNA remained transcriptionally active and was associated with reproducible changes in recipient gene-expression programs and heritable phenotypic variation. Together, our results defined an extrachromosomal route by which graft-mediated somatic exchange can be converted into vertical inheritance, broadening the repertoire of heritable genetic entities beyond chromosomal sequence.

### Periclinal chimeras enable lineage tracing of exogenous eccDNA into sexual progeny

Periclinal chimeras maintain a layered shoot apical meristem (SAM) in which genetically distinct cell layers are spatially organized yet stably maintained, providing a stringent framework to test inter-lineage transfer while preserving cellular boundaries ^11,12^. We used this architecture to trace graft-acquired eccDNA into germline-derived progeny through two independent routes (Fig. 1). In one route, selfed TTC chimeras yielded cp-TTT progeny derived from the L2 lineage after somatic eccDNA exchange. In the other, plants regenerated from the L1 lineage of TCC chimeras were selfed to generate rcp-TTT progeny (Fig. 1; Extended Data Fig. 6). Because these routes differed in lineage provenance but converged genetically, they reduced ambiguity in lineage attribution and provided a robust test of soma-to-germline transmission. Using this system, we found that graft-acquired eccDNA is not confined to somatic layers but can access germline lineages and be detected in sexual progeny across multiple generations (Fig. 3a–c). Progeny carrying these eccDNA displayed stable, heritable phenotypic variation (Fig. 6a–e; Extended Data Fig. 6), linking horizontal somatic exchange to vertical transmission through sexual reproduction. Although these observations establish reproducible graft-induced phenotypic variation in chimera-derived lineages, directly attributing specific traits to individual eccDNAs will require targeted functional perturbation and genetic tests of sufficiency.

### Sequence and structure shape eccDNA stability across the soma–germline boundary

eccDNA transmitted to sexual progeny were found to be enriched for inverted repeats, hairpin-forming sequences, and ACS-like motifs (Fig. 4a–g; Extended Data Tables 16 and 17), suggesting selective retention of architectures compatible with episomal maintenance. ARS-like motifs and structure-forming sequences in plant eccDNA have been reported to support autonomous maintenance under selection, as exemplified by herbicide-resistance eccDNA replicons ^32–34^. Consistent with this enrichment, these cis-encoded features supported autonomous maintenance in yeast (Fig. 5f and 5g). Recent work in other systems further indicated that episomal inheritance can depend on chromatin-associated retention in addition to replication competence; whether analogous retention processes contribute to plant eccDNA inheritance remains to be determined ^65,66^. Transmission patterns were reproducible across both chimera types. Specific eccDNA molecules recurred across independent biological replicates and multiple progeny lineages (Fig. 5a-d). Together, the reproducibility of eccDNA recovery and the enrichment of shared sequence and structural features support a model in which persistence across the soma–germline boundary is biased toward particular eccDNA architectures.

### Transcriptionally active eccDNA may contribute to transgenerational variation

Inherited eccDNA is not uniformly transcriptionally inert. A subset detected in sexual progeny produced transcripts absent from self-grafted controls (Fig. 7e), indicating that some inherited eccDNA remains transcriptionally competent after germline transmission ^26^. Integrating eccDNA profiles with transcriptomics showed a coherent transcriptional response rather than isolated changes, with pathways linked to leaf development, cell-cycle regulation, primary metabolism and drought responses consistently affected (Fig. 7e). Although progeny lines shared a core signature, lineage-specific expression patterns tracked differences in eccDNA composition and abundance, consistent with quantitative and context-dependent effects. Such dose-sensitive regulatory outputs are consistent with observations for extrachromosomal DNAs in other systems ^52,67^. An important next step will be to connect genotype-to-phenotype at the level of individual circles by combining eccDNA engineering or selective depletion with quantitative readouts of eccDNA copy number, eccDNA-derived transcription, and corresponding shifts in the recipient transcriptome and traits.

### A novel extrachromosomal route to heritable variation

The maintenance and transmission of eccDNA across multiple sexual generations, together with reproducible phenotypic effects in progeny, positions eccDNA as a distinct class of heritable genetic elements. Building on our earlier demonstration of graft-induced eccDNA exchange between somatic tissues ^26^, these results define a route in plants in which eccDNA can move between somatic lineages and be transmitted through sexual reproduction. This behavior is distinct from chromosomal variants that require stable genomic integration. Instead, eccDNA comprises discrete extrachromosomal DNA molecules capable of autonomous maintenance and, in some cases, transcriptional activity ^20,21^. This represents a major conceptual advancement in the central dogma of plant genetics by extending the scope of heritable genetic material beyond chromosomal sequence to include extrachromosomal DNA. Collectively, our findings expand the repertoire of heritable genetic entities by introducing a transmissible extrachromosomal genetic component, providing a molecularly defined route by which somatic exchange can contribute to heritable variation ^3^.

### Outlook: from graft interfaces to germline transmission

Grafting is widely used in agriculture for physiological benefits, yet graft interfaces can also enable DNA-level exchange under certain conditions, including transfer of large DNA segments and plastid genomes ^68,69^. In parallel, eccDNA occurs broadly across plant species ^20,21,44^, raising the possibility that soma-to-germline transmission of eccDNA may extend beyond specialized periclinal chimeras and operate with efficiencies shaped by developmental timing and physiological state. It remains unclear how eccDNA moves across tissues, is maintained in germline lineages, and balances episomal persistence and chromosomal integration. Quantitative benchmarks for transmission frequency, stability, and trait predictability across species and environments will be essential to assess the generality and utility of graft-mediated eccDNA transfer. In this context, graft-transmissible eccDNAs provide a concrete, molecularly testable substrate for generating heritable variation without chromosomal integration, with potential relevance to crop improvement.

## Methods

### Plant materials and growth conditions

Periclinal chimeras between tuber mustard (*B. juncea var. tumida*) inbred line and red cabbage (*B. oleracea var. capitata*) were previously generated by in vitro grafting as described . Two types of chimeras were obtained: TCC, in which the shoot apical meristem (SAM) consisted of an L1 layer derived from *B. juncea* and L2–L3 layers from *B. oleracea*; and TTC, in which L1–L2 originated from *B. juncea* and L3 from *B. oleracea*. Self-grafted *B. juncea* and *B. oleracea* plants were designated g-TTT and g-CCC, respectively. Regeneration from the L1 layer of g-TTT and g-CCC produced asexual progenies rg-TTT and rg-CCC^16^, whose selfed offspring were referred to as rgp-TTT and rgp-CCC. In the present study, TCC was used as the starting material. L1 cells of *B. juncea* origin were selectively induced by tissue culture to regenerate whole plants, designated rc-TTT, and their selfed progeny constituted the rcp-TTT1-n population. TTC plants were directly self-pollinated to generate the cp-TTT1-n population. All *Brassica* materials were maintained in vitro at the Laboratory of Germplasm Innovation and Molecular Breeding, Zhejiang University.

Two newly established *B. juncea* germplasm lines, rcp-TTT1 and cp-TTT2, were used as experimental materials. Self-grafted progenies (rgp-TTT and rgp-CCC), together with the parents TTT and CCC, served as controls. Seeds were germinated and transplanted into 50-cell trays containing a substrate mixture of peat, vermiculite and perlite (2:1:1, v/v/v). Plants were grown in a controlled growth chamber at 25 °C under a 16 h light/8 h dark photoperiod. At the four-leaf stage, fully expanded leaves from comparable developmental positions were collected. Leaves from three plants were pooled as one biological sample, and two biological replicates were prepared for each group unless otherwise stated.

### Leaf morphology analysis

Leaf length, width and area were measured to compare leaf morphology among chimeric progenies (rcp-TTT1 and cp-TTT2) and control lines (rgp-TTT and rgp-CCC). Digital images were analyzed using ImageJ software (Fiji; https://imagej.net/software/fiji/). Three biological replicates were analyzed for each group.

### Circle-seq library preparation and sequencing

Six Brassica materials were collected for eccDNA sequencing. For each sample, tissues from three plants were pooled, and two biological replicates were prepared. Genomic DNA was extracted using the CTAB method^70^. eccDNA enrichment was performed following a published Circle-seq protocol with minor modifications. Briefly, eccDNA was isolated using the TIANprep Mini Plasmid Kit (TIANGEN). The extracts were digested overnight with the rare-cutting restriction enzyme MssI (Thermo Scientific) to fragment linear chromosomal and organellar DNA, followed by treatment with Plasmid-Safe ATP-Dependent DNase (Epicentre) to remove residual linear DNA. Enriched circular DNA was amplified using the REPLI-g Midi Kit (QIAGEN). Amplified products were fragmented by ultrasonication, converted into sequencing libraries and subjected to high-throughput sequencing. Circle-seq library construction and sequencing were performed by DIATRE Biotechnology (China).

### Identification and annotation of eccDNAs

Raw paired-end reads were quality filtered (Q30) and processed using fastp (v 0.22.0) to remove adapters and low-quality reads^71^. Clean reads were aligned to the reference genomes of *B. juncea* (http://39.100.233.196:82/download_genome/Brassica_Genome_data/Braju_tum_V2.0/) and *B. oleracea* (https://db.cngb.org/data_resources/assembly/CNA0003257/) using BWA (v0.7.12)^72^. eccDNAs were identified with Circle-Map (v1.1.4) ^73^.

Soft-clipped reads at eccDNA breakpoints were extracted using SAMtools (v1.4.1) (http://samtools.sourceforge.net). eccDNAs were annotated using BedTools (v2.29.2) ^74^. Differential expression of ecGenes was identified based on normalized junction read counts using edgeR (fold change ≥ 2.0, *P* ≤ 0.05), followed by gene-level annotation, and enrichment analysis ^42^. Visualization of eccDNA structures was performed using IGV (v2.4.10). To investigate the presence of conserved sequences in eccDNAs maintained in sexual offspring, the sequence features of eccDNAs were examined using MEME (https://meme-suite.org/meme/opal-jobs/appMEME_5.5.717255067415341578431199/meme.html#sites_sec).

### Direct and inverted repeat analysis of eccDNAs

To analyze the sequence characteristics of eccDNA, we extended the end coordinates of eccDNA by 150bp outward and contracted them by 150 bp inward and used short blastn to identify inverted and direct repeat sequences in the resulting sequence.

### Inverse PCR validation of eccDNAs

Genomic DNA was extracted using the CTAB^70^ method and treated with Plasmid-Safe ATP-Dependent DNase (Lucigen) for 128 h to remove linear DNA. Rolling-circle amplification (rRCA) was then performed, and products were purified using the Genomic DNA Clean &

Concentrator™-10 kit (Zymo Research). Untreated genomic DNA, DNase-treated eccDNA and rRCA products were used as templates for inverse PCR.

Inverse primers were designed using NCBI BLAST (https://blast.ncbi.nlm.nih.gov/Blast.cgi), and are listed in Extended Data Table 20. PCR amplification was performed using either Phanta Max Master Mix (Vazyme) or Taq DNA Polymerase with ThermoPol® Buffer (NEB). PCR products were resolved by agarose gel electrophoresis, purified using the E.Z.N.A.® Gel Extraction Kit (Omega), and validated by Sanger sequencing, Illumina-based KBseq or Oxford Nanopore sequencing. Sequence visualization was performed using karyoploteR and DNAMAN.

### Sequence analysis of DNA A + T content and curvature

Genomic A + T and G + C content was globally determined for the eccDNA replicon by dividing the sequence into 50 bp sequential windows with the MakeWindows function of BedTools v.2.29.2^74^. A + T and G + C content was determined for each window using the nuc function of BedTools v2.29.2^74^ and plotted as a circular track using the Circos v.0.69.9^75^. DNA curvature analysis was performed by extracting a 240 bp and 397 bp segment from the eccDNA replicon sequence (coordinates: C5:42317656–42319060) and analyzed for curvature using the online version of dnacurve 2020.1.1 (17 Jan, 2020 release) (https://pypi.org/project/dnacurve/) using the wedge model to determine the axial path of this segment of DNA. The resulting protein data bank (PDB) file was visualized with the UCSF Chimera tool^76^.

### Cloning and functional verification of eccDNA autonomous replication sequences (ARS) in yeast

The full-length eccDNA sequence corresponding to C5:42317656–42319060, as determined by first-generation (Sanger) sequencing, was submitted to Tubegene for de novo synthesis of a circular DNA molecule. The eccDNA-derived ARS-containing fragments and inverted repeats and hairpin structures were subsequently amplified using the synthetic eccDNA as the template with primers eccARS-F and eccARS-R (Extended Data Table 20). The yeast vector, pRS315, was linearized via PCR using primers pRS315ΔARS-F and pRS315ΔARS-R such that the CEN6 sequence remained, but the native ARS was removed. Q5 polymerase was used for all PCRs. The eccDNA ARS region (eccARS) was assembled into pRS305 and the ARS-less pRS315 using a sequence and ligation independent cloning (SLIC) reaction^77^. Constructs were confirmed with a restriction digest and sequencing. Saccharomyces cerevisiae (ATCC 208288) were transformed in triplicate as previously described. Yeast cells were grown in a YPD (10 g/L yeast extract, 20 g/L peptone, 20 g/L glucose) preculture overnight at 28 °C and 250 rpm. In a 250-mL baffled flask, 50 mL of pre-warmed YPD was inoculated to a final titer of 5 × 106 cells/mL. The culture was grown to a final titer of 2 × 107 cells/mL at 28 °C and 250 rpm. Cells were harvested by centrifugation at 3000×g for 5 min. The cell pellet was resuspended in 25 mL of sterile milli-Q water and centrifuged again three times before cells were resuspended in 1.0 mL of sterile water. Cell pellet was then resuspended in 360 μL freshly made transformation mix (240 μL PEG 3350 (50% w/v), 34 μL 1.0 M LiAc, 50 μL single-stranded salmon sperm DNA (2 mg/mL), 36 μL plasmid DNA plus sterile water). Cells were heat shocked at 42 °C for 40 min and resuspended in 1 mL of sterile milli-Q water. 200 μL of cells were plated on YSC Leu + 2% glucose plates and grown at 28 °C for 2 days and colonies counted. pRS305 lacks an ARS and served as a negative control while pRS315 contains an ARS and served as a positive control. To confirm plasmid retention, colonies from trans formed plates were passaged on YSC-Leu + 2% glucose plates three times. The passaged cells were used for a col ony PCR using Q5 polymerase and primers eccARS-F/ eccARS+hairpin-F and eccARS-R/ eccARS+hairpin-R. The original colonies from the pRS305 transformation plate were used as templates, as they did not survive passaging. Positive controls using plasmids pRS305+eccARS/ pRS305+eccARS + hairpin and pRS315+eccARS+hairpin and a negative control using wild type *S. cerevisiae* cells were performed. The PCR products were run on a 1% TAE gel with Gen script Ready-to-Use™ Plus 100 bp DNA Ladder.

### RNA sequencing and transcriptome analysis

Total RNA was extracted from fully expanded leaves at the four-leaf stage using TRIzol reagent (Invitrogen). Two biological replicates were prepared for each treatment. RNA-seq libraries were constructed using the NEBNext Ultra RNA Library Prep Kit for Illumina and sequenced on the NovaSeq 6000 platform to generate 150-bp paired-end reads.

Adapters and low-quality reads were removed using fastp (v0.23.2)^71^. Clean reads were aligned to the *B. juncea* (TTT) reference genome (Index of/download_genome/Brassica_Genome_data/Braju_tum_V2.0) using HISAT2 (v2.0.5) (Extended Data Table 21) ^78^. SAM files were converted to BAM format with SAMtools (v1.4.1) (http://samtools.sourceforge.net). Transcript assembly was performed using StringTie^78^, and gene expression was quantified with FeatureCounts^79^. Differential expression analysis was conducted using DESeq2^80^ (fold change > 1.5, adjusted *P* < 0.05). GO and KEGG enrichment analyses were performed using ClusterProfiler.

### Drought stress treatment

Drought stress was imposed by withholding water for 0, 3 and 7 days. Plants maintained under normal watering conditions served as controls.

### Physiological measurements

The Brassica plants were photographed with a Canon EOS 80D to obtain high-resolution images. 0, 3 and 7 days after treatments, leaves were harvested and weighted immediately after detached from plants. Relative moisture content (RWC) was detected on 15 leaves for each treatment. Fresh weight (FW) was recorded and then leaves were placed in distilled water at 4 °C for 24 h to get saturated weight (SW). Finally, leaves were dried at 65 °C for 6 h to determine dry weight (DW). RWC was calculated as the following formula^81^: RWC = [(FW — DW) / (SW — DW)] × 100% 0.3 g leaf disks (0.6 cm of diameter) from each treatment were harvested for relative electric conductivity (REC) measurements. Samples were placed in a 50 mL tube containing 30 mL distilled water and sharked at 200 rpm at 28 °C for 2 h, then the initial electrical conductivity (K1) was measured with a conductivity meter. After that, the tubes were boiled at 95 °C for 20 min then cooled to room temperature. Finally, the final electrical conductivity (K2) was measured. REC was calculated as the following formula ^82^: REC = K1/K2 × 100%

### Microscopic observation

For light microscopy observation, the samples were fixed in a formaldehyde: acetic acid: ethanol fixative, washed twice with 70% v/v ethanol, and dehydrated in a graded ethanol series at room temperature (70%, 85%, 90%, and 95% v/v for 50 min, followed by 100% for 30 min, twice). The samples were infiltrated with a mixture of pure xylene and pure alcohol (1:1) for 30 min and subsequently infiltrated with pure xylene twice for 30 min. Shredded paraffin was slowly poured onto the samples containing xylene at room temperature to achieve a 50% v/v ratio of paraffin: xylene; the paraffin-embedded samples were then incubated overnight at 37℃. On the morning of the next day, the paraffin-immersed vessel was placed in an incubator at 60℃, the caps of the tubes were opened, and xylene was allowed to volatilize for 2 h. The samples were then transferred to fresh 100% paraffin in an incubator at 62 °C for 6 h, and the paraffin was replaced every 2 h. The pre-melted (60 ℃) pure paraffin was quickly poured into the embedding box. Once paraffin at the bottom of the embedding box solidified, the samples were moved rapidly into the embedding box, which was filled with paraffin. For light microscopy, polymerized samples were sectioned (8 μm thickness) with a paraffin slicing machine and mounted on glass slides. For light microscopy observation, the sections were stained with 0.5% w/v toluidine blue (pH 7.0).

## Acknowledgements

We are grateful to Dr. Zhenyu Qi and Mr. Rong Jin for providing facilities and assistance with plant material management at the Agricultural Experiment Station of Zhejiang University; to Dr. Mingfang Zhang and Dr. Ting Zhao for valuable suggestions.

## Author contributions

Liping Chen and Aijun Zhang conceived and designed this research. Liping Chen performed the construction of artificial Brassica chimeras. Aijun Zhang, Han Zhou, and Ke Liu maintained the chimeric plants and their progeny. Liping Chen, Aijun Zhang, Han Zhou, and Ke Liu conducted regular field observations. Aijun Zhang, Han Zhou, Kexin Xu, Tingjin Wang, Junxing Li, Lu Yuan, Li Huang, Yongfeng Guo, Yan Wang and Ningning Yu performed the experiments. Aijun Zhang, Han Zhou and Wanmu Zhou designed the graphical pictures and diagrams. Aijun Zhang and Liping Chen wrote the manuscript. Yanhong Zhou, Susheng Gan, Hannah Rae Thomas, Ke Liu and Liping Chen revised the manuscript. Aijun Zhang, Han Zhou, Wanmu Zhou and Qinjie Chu analyzed the data. All authors read and approved the manuscript.

## Data availability

The raw sequencing data have been deposited in GSA (Genome Sequence Archive in BIG Data Center, Beijing Institute of Genomics, Chinese Academy of Sciences, http://gsa.big.ac.cn) under the accession number CRA038188. All the other data supporting the findings of this study are available within the article and its supplementary information files and from the corresponding author upon reasonable request. Source data are provided with this paper.

## Ethics declarations

### Ethics approval and consent to participate

Not applicable.

### Consent for publication

All authors have read and approved the final version of the manuscript.

## Extended data figures and tables

**Extended data Fig. 1.**
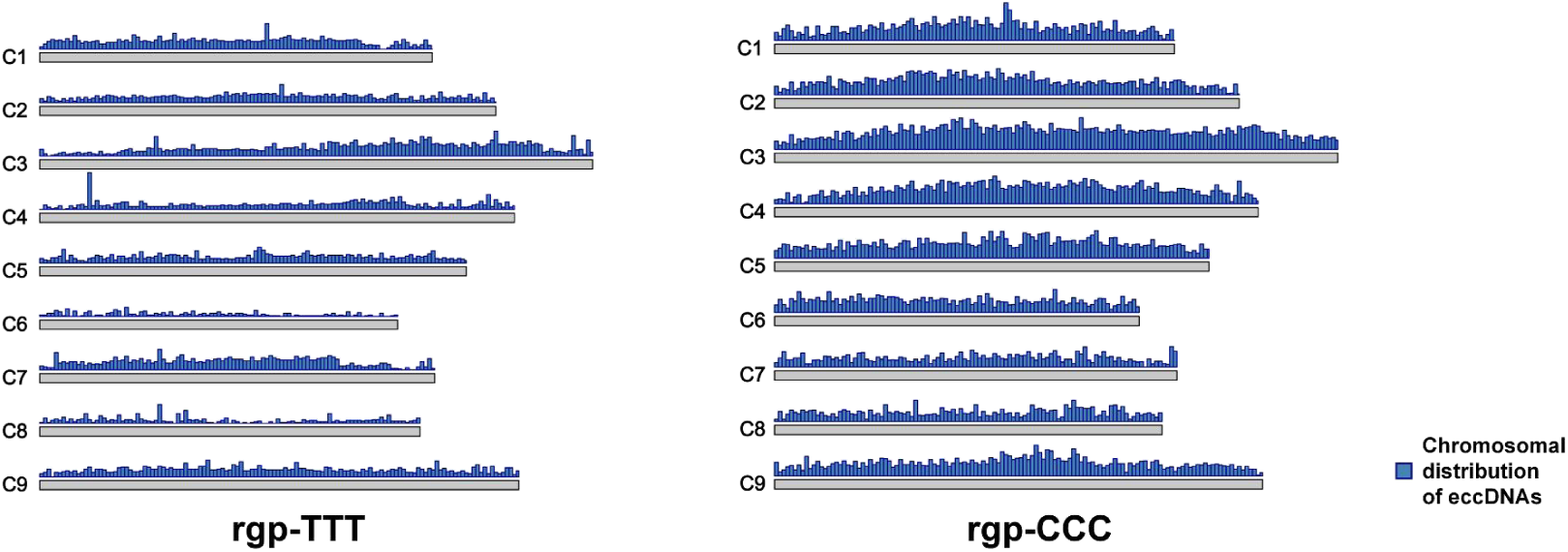
Karyoplot displaying the chromosomal distribution of *B. oleracea* eccDNAs found in rgp-TTT and rgp-CCC.

**Extended data Fig.2.**
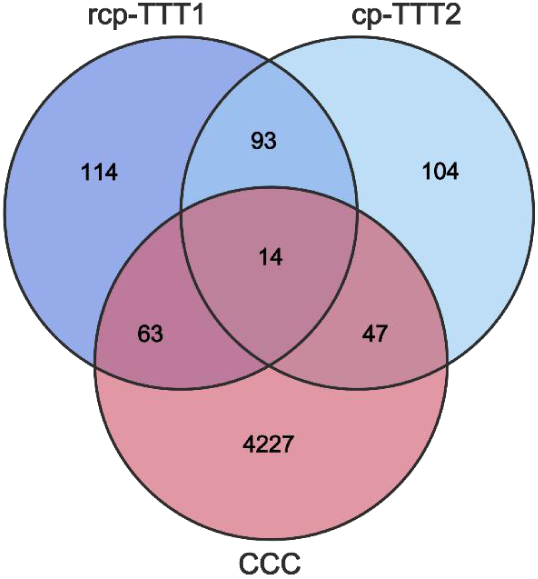
Venn diagrams displaying the overlaps of transferred *B. oleracea-*specific eccDNAs relative to the red cabbage genome across different Brassica tissues in rcp-TTT1, cp-TTT2, and CCC.

**Extended data Fig. 3.**
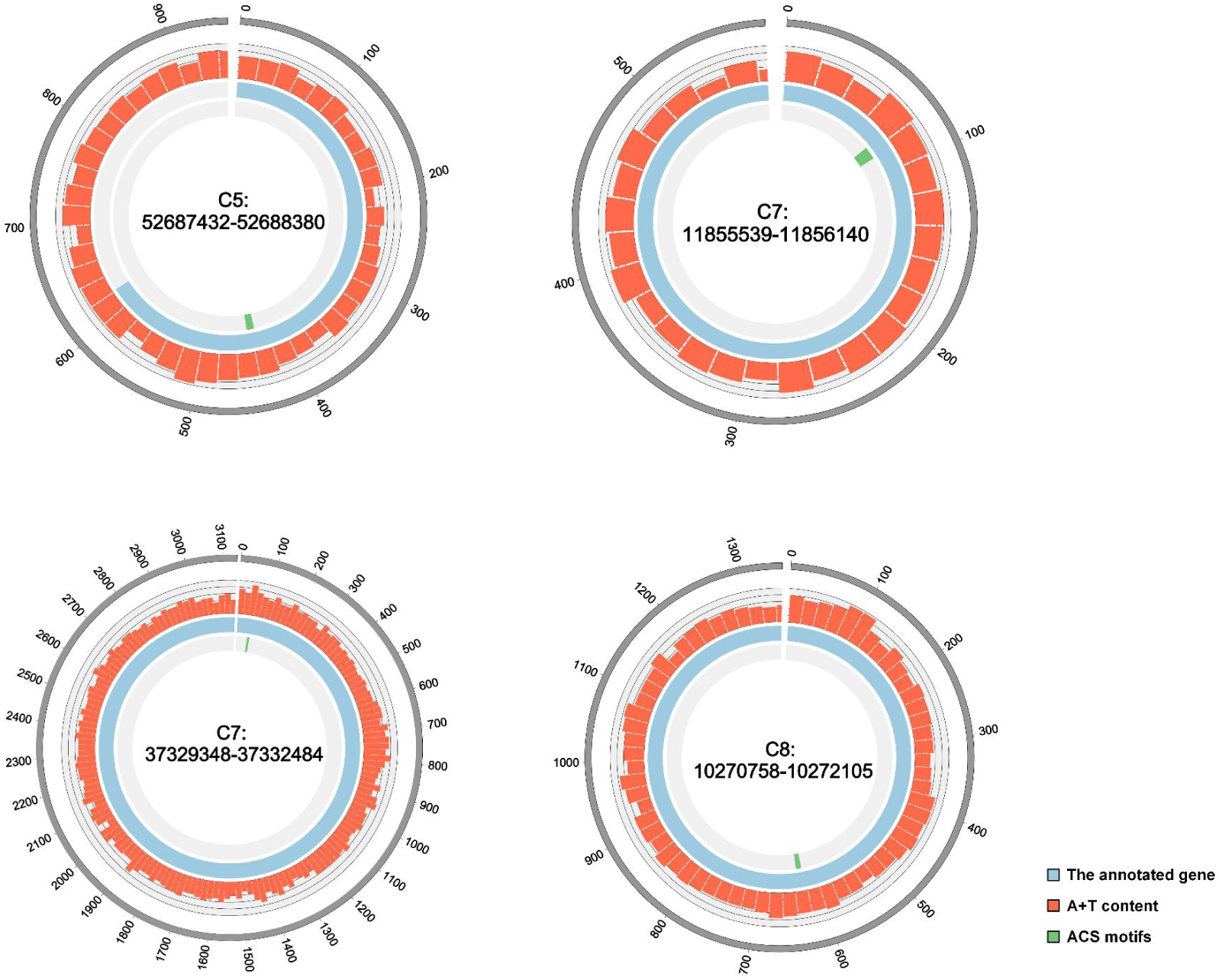
Circular map of the eccDNA replicons C5: 52687432-52688380, C7: 11855539-11856104, C7: 37329348-37332484 and C8:10270758-10272105 in rcp-TTT1. The A+T content distribution was calculated using 50-bp sliding windows. The outermost ring of the map indicates the genomic origin of the eccDNA; the red track represents the A+T content; blue blocks correspond to the annotated gene *Boc05g03996* in C5: 52687432-52688380, *Boc07g01715* in C7: 11855539-11856104, *Boc07g03746* in C7: 37329348-37332484, *Boc08g01595* in C8:10270758-10272105; and the innermost green track marks the positions of ACS motifs.

**Extended data Fig. 4.**
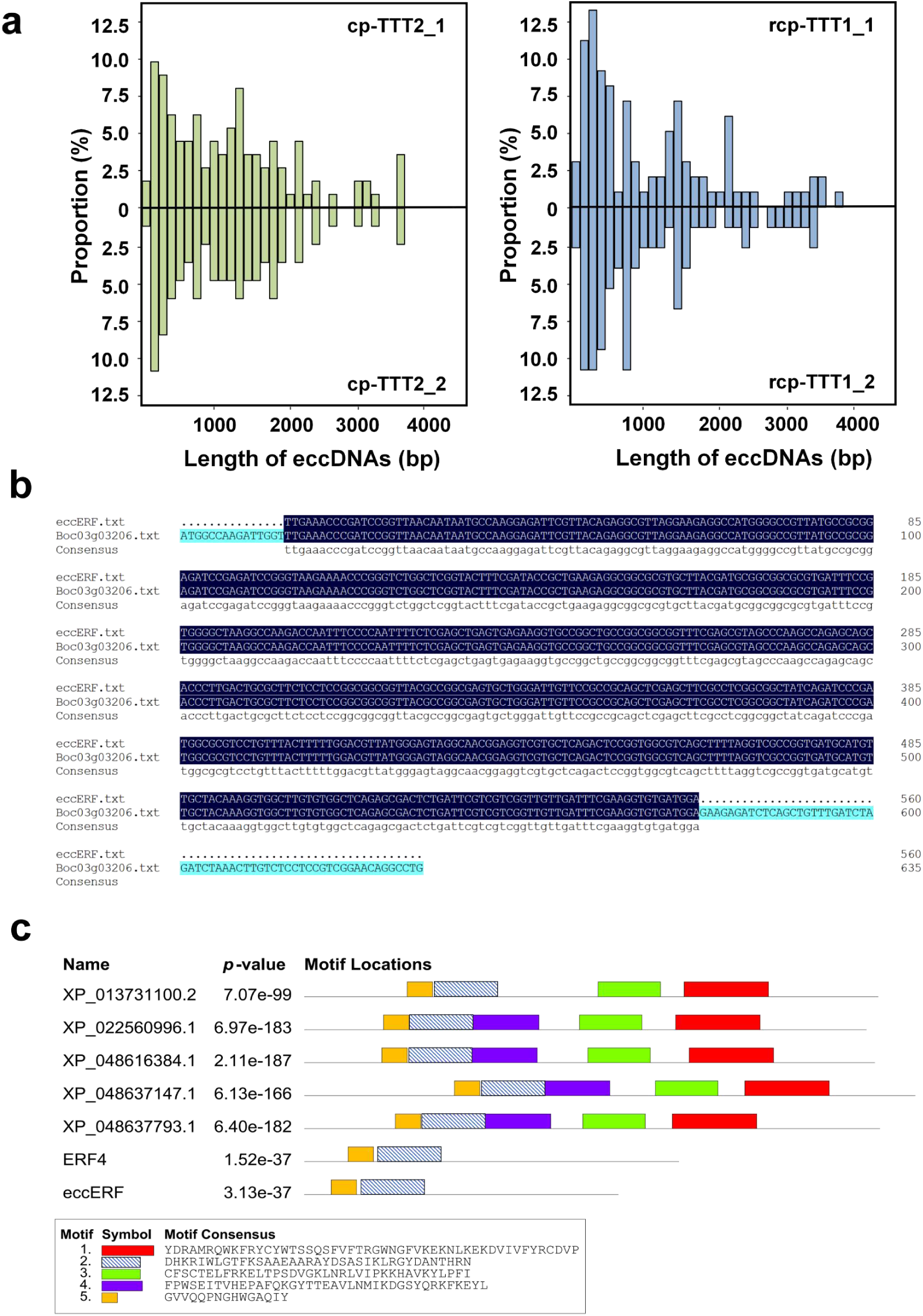
**a,** Length distribution of eccDNAs in cp-TTT2 and rcp-TTT1 (two biological replicates each), ranging from 139 bp (minimum) to 5,212 bp (maximum). **b,** The sequence alignment similarity between *eccERF* and *Boc03g03206.* **c,** Conserved sequence elements identified using the MEME suite in transferred eccDNAs detected in the sexual progeny. Red, purple, and green symbols represent motifs identified in homologs of the *ERF4* gene family.

**Extended data Fig. 5.**
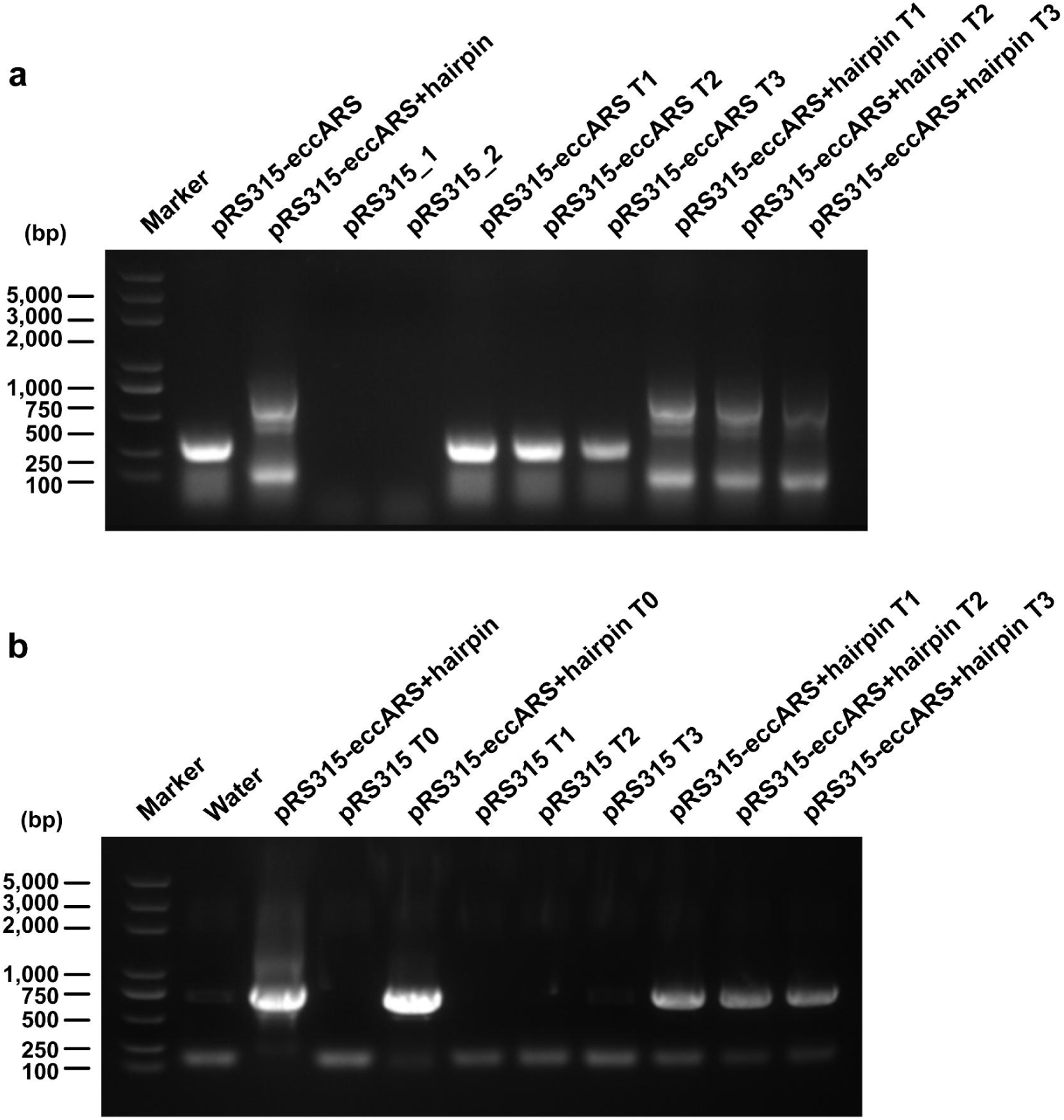
Gel results for a colony PCR. Lanes are as follows: L, ladder; **a,** 1, pRS315ΔARS + eccARS + CEN6 plasmid; 2, pRS315ΔARS + eccARS + hairpin + CEN6 plasmid; 3–4, pRS315 replicating plasmid; 5–7, triplicates of passaged pRS315ΔARS + eccARS + CEN6 transformed cells; 8–10, triplicates of passaged pRS315ΔARS + eccARS + hairpin + CEN6 transformed cells. **b,** 1, water; 2, pRS315ΔARS + eccARS + hairpin + CEN6 plasmid; 3, T0 of pRS315 replicating plasmid; 4, T0 of pRSpRS315ΔARS + eccARS + hairpin + CEN6 plasmid; 5–7, triplicates of passaged pRS315 replicating plasmid; 8–10, triplicates of passaged pRS315ΔARS + eccARS + hairpin + CEN6 transformed cells.

**Extended data Fig. 6.**
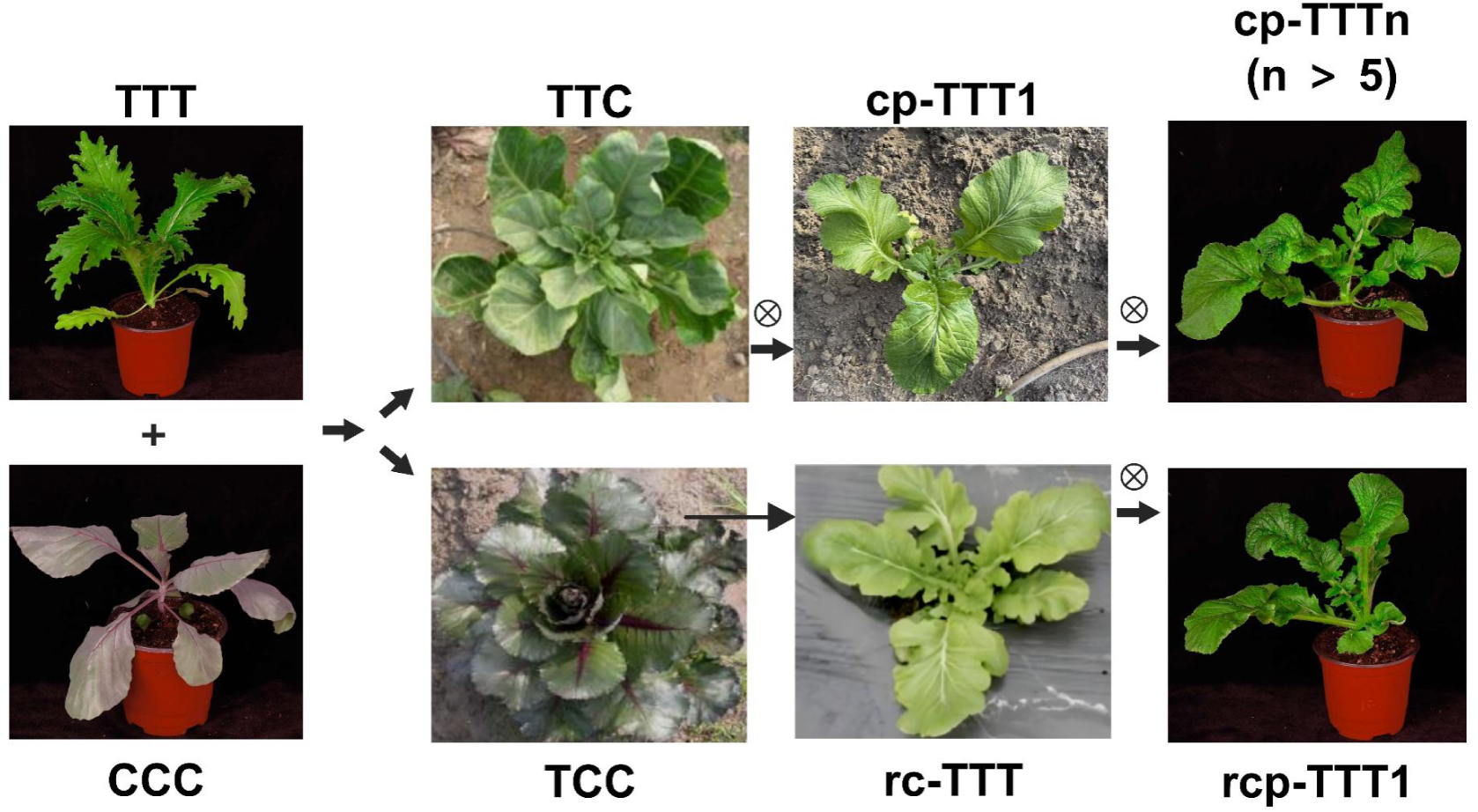
Phenotypes of Brassica parents and graft chimeric materials, along with their sexual progenies.

**Extended data Fig. 7.**
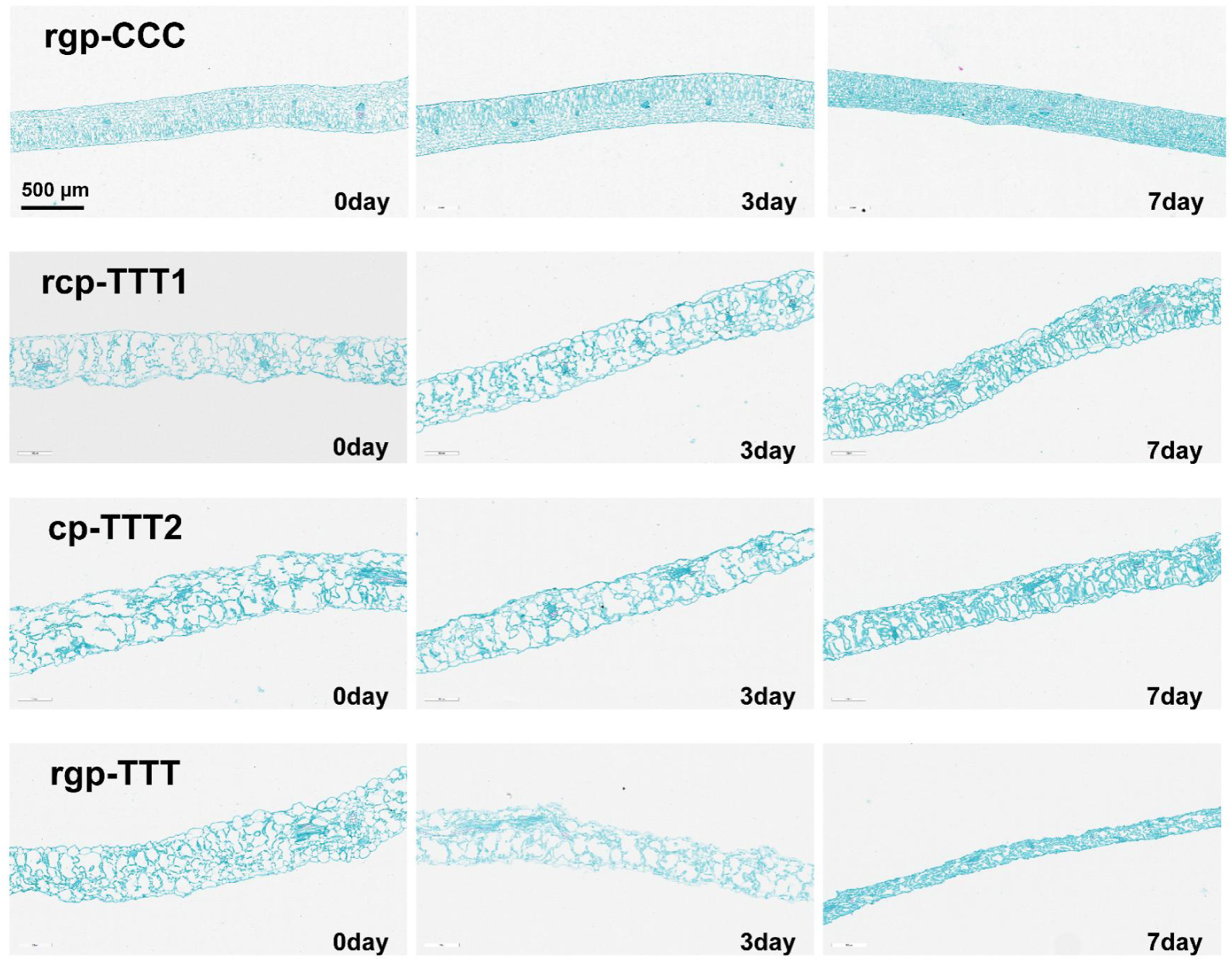
Leaf paraffin sections of rgp-CCC, rcp-TTT1, cp-TTT2, and rgp-TTT following different durations of drought treatment.

**Extended data Fig. 8.**
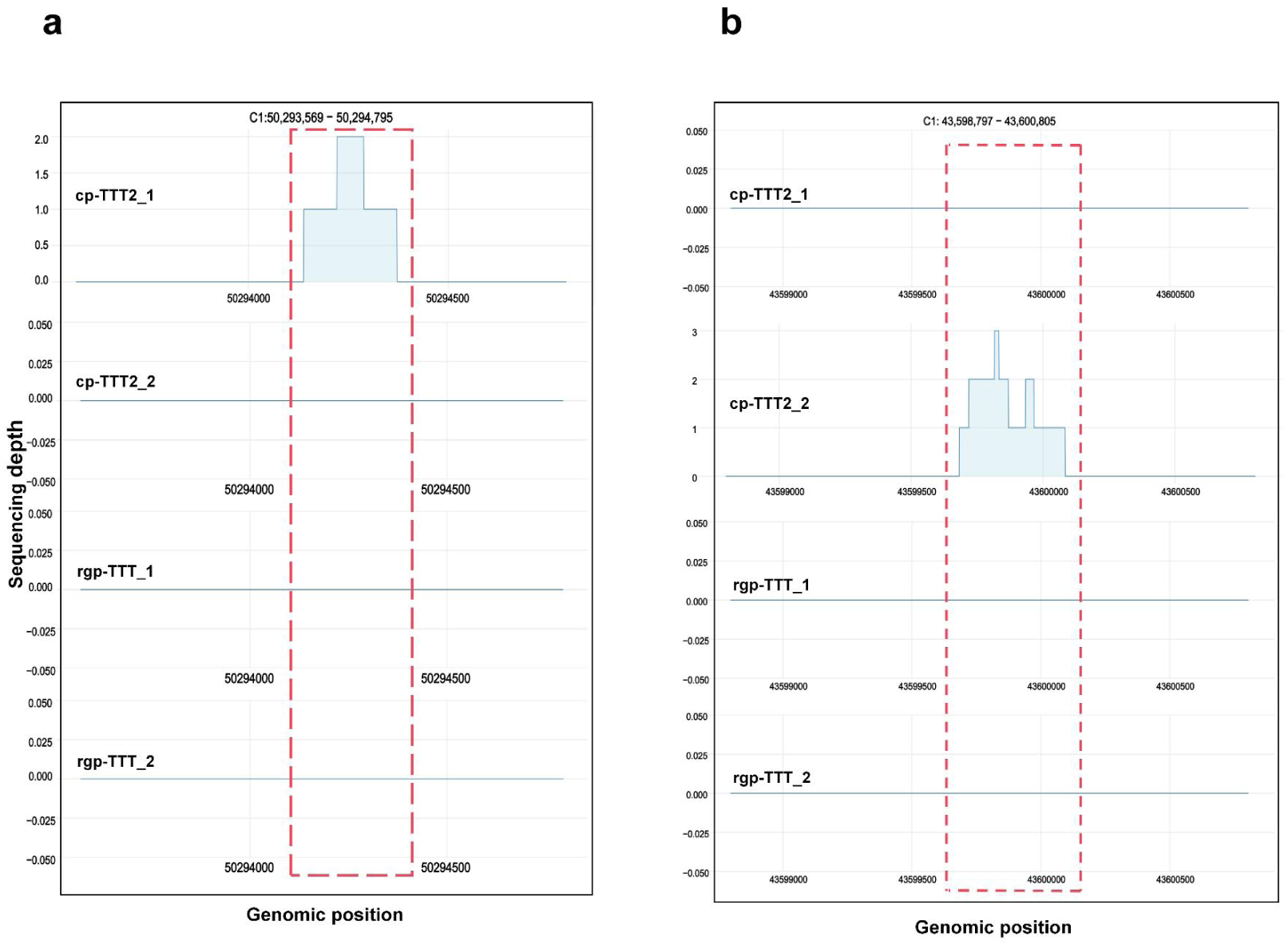
Integration of transcriptome and eccDNA mapping data in cp-TTT2 (cp-TTT21 and cp-TTT22) and rgp-TTT (rgp-TTT11 and rgp-TTT12).

**Extended data Fig. 9.**
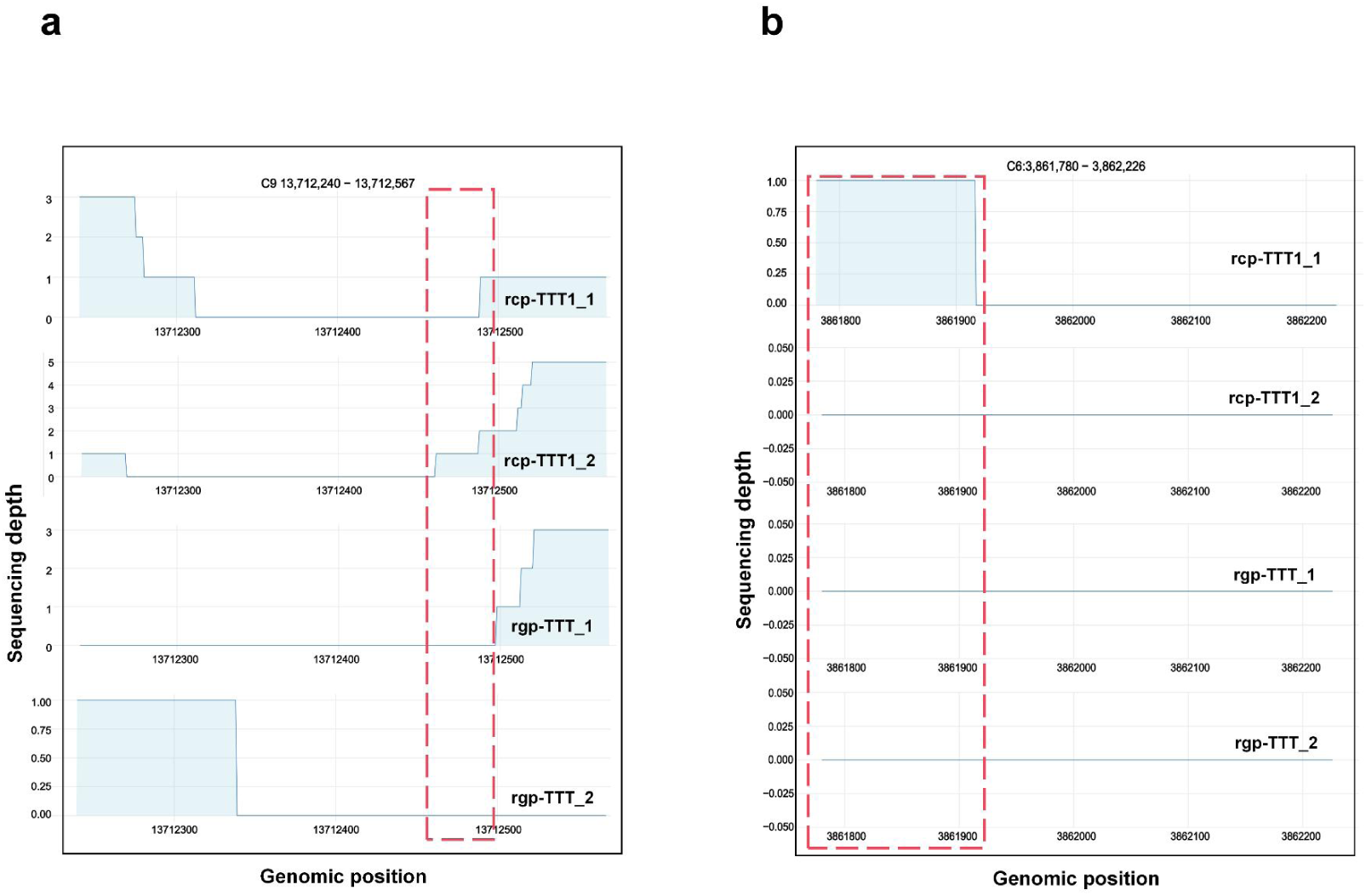
Integration of transcriptome and eccDNA mapping data in rcp-TTT1 (rcp-TTT11 and rcp-TTT12) and rgp-TTT (rgp-TTT11 and rgp-TTT12).

**Extended Data Table 1. eccDNA data-filtering statistics.**

**Extended Data Table 2. Reads uniquely aligned to the *B. oleracea* reference genome.**

**Extended Data Table 3. eccDNAs identified by Circle-Map.**

**Extended Data Table 4. Transferred *B. oleracea* – specific eccDNAs detected in two sexual progenies.**

**Extended Data Table 5. Length distribution of eccDNAs in cp-TTT2 and rcp-TTT1.**

**Extended Data Table 6. Median GC content of eccDNAs in cp-TTT2 and rcp-TTT1 compared with genomic regions 150 bp upstream and downstream of their source loci.**

**Extended Data Table 7. Transferred eccDNAs with inverse repeats in cp-TTT2 and rcp-TTT1.**

**Extended Data Table 8. Transferred eccDNAs with direct repeats in cp-TTT2 and rcp-TTT1.**

**Extended Data Table 9. Transferred eccDNAs overlapping with gene regions and intergenic region.**

**Extended Data Table 10. Transferred eccDNAs overlapping with exons, introns, upstream 2 kb regions (up2kb) and downstream 2 kb regions (down2kb).**

**Extended Data Table 11. Count of transferred eccDNAs associated with a singular gene, and eccDNA spanning two genes in cp-TTT2 and rcp-TTT1.**

**Extended Data Table 12. eccDNAs containing genes (GcE) and genes containing eccDNAs (EcG) transferred in cp-TTT2 and rcp-TTT1.**

**Extended Data Table 13. Conserved ACS-like 11-bp motifs were identified in eccDNAs transferred into rcp-TTT1 and cp-TTT2.**

**Extended Data Table 14. ACS motifs were located within genes.**

**Extended Data Table 15. AT content of the ACS motifs, 50 bp up and down within the ACS motifs and further across entire eccDNAs in rcp-TTT1 and cp-TTT2.**

**Extended Data Table 16. Hairpin structures were identified in transferred eccDNAs in rcp-TTT1 and cp-TTT2.**

**Extended Data Table 17. Hairpin arms directly spanned the 11-bp core sequence.**

**Extended Data Table 18 Information of *B. oleracea*–specific transferred eccDNAs detected in the two sexual progenies, cp-TTT2 and rcp-TTT1, each with two biological replicates.**

**Extended Data Table 19. Read abundance of transferred eccDNAs different sexual progeny with two replicates.**

**Extended Data Table 20. Primer sequences for PCR confirmation of eccDNA presence.**

**Extended Data Table 21. Transcriptome data filtering statistics.**

**Extended Data Table 22. Unique transcripts of cp-TTT2 and rcp-TTT1 compared with transferred eccDNAs.**

## Supplementary information

C3:46317085–46317645. Sanger sequencing. Txt

C5:42317656–42319060. Sanger sequencing. txt

