## Supplemental Data 1 for "Somatic-to-germline transmission of horizontally acquired extrachromosomal circular DNA in Brassica graft chimeras"

C3：46317085﹣46317645

GCAGCACCCTTGACTGCGCTTCTCCTCCGGCGGCGGTTACGCCGGCGAGTGCTGGGATTGTTCCGCCGCAG
CTCGAGCTTCGCCTCGGCGGCTATCAGATCCCGATGGCGCGTCCTGTTTACTTTTTGGACGTTATGGGAGTA
GGCAACGGAGGTCGTGCTCAGCCTCCGGTGACGTCAGCTTTTATGTCGCCGGTGATGCATGTGGCTACAAA
GGTGGCTTGTGTGGTGGCTCAGAGTGACTCTGATTCGTCTTCGGTTGTTGATTTTGAAGGCGTGATGGAGA
AGAGATCTCAGCCGTTAGATCTAGATCTAAACTTGCCTCCTCCGTCGGAACAGGCCTGAACTGATCGAAGGT
GGTCGTGGTCGTGTTCACATCCGACGCGCATGCGTTTTTTTTTTTTTTTGTTTGGGAGCTGAGAAAAAAAAC
TTTTTTCTCGTAGTTTTTATTTCTCTCACTCAACAATAATTATTATCGGTTTTAAAAAATTAAATCGGGAGAGAA
GAAAATAATTTTAGCTGACATGGCGTGCTTACGACGCGGCGGCGCGTGATTTCCGCGGGGCCAAGGCCAA
GACCAATTTCCCCAATTTTCTCGAGCTGAGTGAGAAGGTGCCGGCTGCCGGCGGCGGTTTCGAGCGTAGCC
CAAGCCAGA
