## Supplemental Data 2 for "Somatic-to-germline transmission of horizontally acquired extrachromosomal circular DNA in Brassica graft chimeras"

C5：42317656－42319060

AGTTCTCTCCCCATGCCTCTCGCCTAGGACTCCTAATATACTCCCCCCTAGGCCGGTTTACGCTTTTCCCCTTCT
GCCCTTAAGGGGTCATAGCCTAAATATGGAGATCTTCCATTGCTTCCTGCTCTATGTTGTAAAGTTATGGTTATA
TTATATATTTCCTAAATAGTATACAAACAACGAAATTACAATATGAATAAGAAATAGAGCTATACTATAAAAATGT
AAAATAGAGTATAGATTTAAAACTTAGTTTATGTAATATAAATTAAATTTATAGAAATATATATGTTTTTAATTTTC
AAAACATATTATGTAGTAAAAATCTATGTTTTTATTATGTAAAATATAAAACTGAATATTTATATATCGGGATTTG
AAAGAGAACACAAAGGAGGATCTTTTTGATGCCTGACAGGTCGCTACATAGCGAGTAGAAGCAAACCAAG
AAGAGTCCTACTTGTTTTCGTCGTAATTTCTCAATGGAAATCCGATTGAGACGTAACGAACAGCGTTTCGATG
GGGATTCGAAAGAGAACGCAAAGGACGACCTTTCTGAGGCCTGACAGGTCGCTATGTAGCGAGTGGAGGT
CTGACAGTTTGCTACGTAGCAAGTAGAAGCTAGCCAAGAAGAGTCCTACTTGTTTTCGTCGTAAAATCTCAA
CGGAAACTCCGATTGAGACGAAATGAAAAGCGTTTCGATGAGGATTCAAAAGAGAACGCAAAGGAGGACT
GACAGGTCGCTATGTAGCGAGTGAAGAGTTGTCTTGAGCTCGGTCGCTACGTAGCGACCGAGCTTAGCTAG
AGCTCGGTCGCTACGTAGTGACCAAGCTGCGTAATCGATTTGCTGTGCTTCCTCTTTCCGCGATTAACCTAGT
TGTTTTCTACGGTTTTTGGGAGAACAAGTTTTAACCTTCAGAAAAGTTTTCGGAAAACGTGTTTTGGTAAAA
TCCTTACGCATTAAGACTTCTTTAGCTCAGAAATGAGCCAAACTTACGGGATTTGATAAAATTTATCGTTTTCC
TTTATGTTTAGACAGAGAAATCGCGAGCTCGTTTCGTAGCACTTCCAGTGGCGAAAAGTTGCATCAAGGTTT
TCGATTTTCTCAAGAACTGTGGCTTTTGCGTAGGCGTTGGCCAAAGGTGCAGTGGCAGCTGGGGTTGTCCT
ACCAGCGTTATTGCGAGCTGCGATTTTGTCTTTCGTATTTCCAGACATTGTTTTGATCAAGCGTTTTGTGGCT
GAGTTAGATGTATCCGTACCCCCCTCCTTCTAGCGCCAAACTGTGGGAACCGAAATTCGCACTGTCGATTTCC
GTTTAAATAAGGAAAGTAGGAGAGCCCTAATTTCCCAGAGGTCCCGGATATCTTCTATTTCCACACGCCAAG
CAATCAGAACACGAAATAAAGATGATAAGATATAAGAAATCGTAAAAAGAGAGCAAAGTAGATCTTATTCCG
AATCTGCGTTTGAGCGTTACAGCAAGGTAAAAGCCTGGGCTACGAGAGTTGTCGGCGAGATTCCTAGTTCT
AAAACCCTAAGACGGCAACAACCTAATTGAGTCGCAGCTCGAATAACAAAAAACAAAAAAATTGCCTAAAC
TGCCCTAAGTGCTAAGTTTGATGTGTAA
